# Evolution of intrinsic disorder in the structural domains of viral and cellular proteomes

**DOI:** 10.1101/2024.10.30.621179

**Authors:** Fizza Mughal, Gustavo Caetano-Anollés

## Abstract

Intrinsically disordered regions are flexible regions that complement the typical structured regions of proteins. Little is known however about their evolution. Here we leverage a comparative and evolutionary genomics approach to analyze intrinsic disorder in the structural domains of thousands of proteomes. Our analysis revealed that viral and cellular proteomes employ similar strategies to increase disorder but achieve different goals. Viral proteomes evolve disorder for economy of genomic material and multifunctionality. On the other hand, cellular proteomes evolve disorder to advance functionality with increasing genomic complexity. Remarkably, phylogenomic analysis of intrinsic disorder showed that ancient domains were ordered and that disorder evolved as a benefit acquired later in evolution. Evolutionary chronologies of domains indexed with disorder levels and distributions across Archaea, Bacteria, Eukarya and viruses revealed six evolutionary phases, the oldest two harboring only ordered and moderate disorder domains and delimiting two universal common ancestors of life. A biphasic spectrum of disorder versus proteome makeup captured the dichotomy in the evolutionary trajectories of viral and cellular ancestors, one following reductive evolution driven by viral spread of molecular wealth and the other following expansive evolutionary trends to advance functionality through massive domain-forming co-option of disordered loop regions.

**Teaser:** Comparative and evolutionary genomic analyses of the intrinsic disorder of structural domains in thousands of proteomes reveal proteomes increase disorder as a benefit acquired later in evolution to support viral economy and multifunctionality or advanced cellular functionality.

## Introduction

Intrinsically disordered regions are flexible regions of proteins that lack a fixed three-dimensional structure under native cellular conditions. These regions are present across the proteins of all organisms in the superkingdoms of cellular life and the proteins of viruses [1]. They possess diverse biophysical properties that allow them to perform a multitude of functions that involve complex signaling and regulation interactions in proteins and nucleic acids. The absence of fixed structure in intrinsically disordered regions that are capable of an assortment of biological functions challenges the long held “one sequence-one structure-one function” paradigm and puts forth the idea of a protein structure-function continuum that adequately explains structure-function relationships in proteins [1,2]. The structure-function continuum model elaborates upon the fact that complexity of biological systems depend on the size of their proteomes but not on the size of their genomes [3].

The dynamic behavior of disordered regions is evolutionarily conserved despite minimal sequence conservation [4]. Flexibility of proteins is also a conserved evolutionary trait, and therefore, measuring disorder can serve as a proxy for the inherent flexibility of a protein [5]. In fact, the flexibility provided by intrinsic disorder in multicellular organisms appears to equip them with a “toolkit” of complex functions such as signaling, regulation and modulating enzyme activity [6]. Disordered regions tend to have, though not necessarily always, a higher evolutionary rate than ordered proteins [7]. However, disordered proteins evolve in a different manner to conserve disorder over different evolutionary timescales.

Intrinsic disorder has been studied extensively at the whole proteome level [8–10]. However, the scope of these studies has been limited to interactions within and not across levels of complexity with little focus on the evolutionary underpinnings of these interactions. Moreover, studies of evolutionary origins of intrinsic disorder have relied on deductions from sequence information [11]. Traditional phylogenetics and sequence alignment-based methods are often limited by the effects of high mutation rates, horizontal gene transfer (HGT), and genetic mosaicism, especially in viral proteomes [12]. The highly conserved nature of protein structure has proven helpful in studying similarities in viruses despite their highly variable sequence makeup, even when sequence similarity is low [13], thereby making them suitable to study similarities in cellular proteomes as well.

In particular, protein structural domains are evolutionarily conserved units of structure and function that harbor deep phylogenomic information [14–16]. The distribution of protein domains across organisms dissects patterns of structural recruitment among superkingdoms, with high levels of explanatory power. Using domains to construct phylogenomic trees and build chronologies [17] has enhanced understanding of the evolution and diversification of cellular life [18] and the origin of viruses [19–21]. Chronologies of domains provide insight into how metals have been utilized by the ‘metallomes’ of prokaryotes and eukaryotes [22]. Tracing domain history in ribosomes uncovered their emergence from primitive ribonucleoprotein entities [23]. Domain chronologies also shed light on the early evolution of modern biochemistry [24], including extensive recruitment occurring in evolving metabolic networks [25–29]. Tree reconstruction using other unconventional genomic characters, such as annotations of molecular function, yield phylogenies that successfully capture the tripartite world of cellular life and many details of organismal diversification [30].

In previous studies, we found that folding speed, which is known to correlate with protein flexibility, decreases and then increases in evolution [31]. This biphasic pattern is puzzling and has been linked to the emergence of vestibular co-translation folding in the ribosome [17]. Investigating patterns of disorder and by extension patterns of flexibility in the structure of proteins can provide important insights into the interplay of protein structure, function, and molecular flexibility. Here we investigate the evolution of intrinsic disorder in biological systems. We focus on protein domains and their relationship with disorder at the proteome level. The interactions within and across these levels of complexity gain support from a phylogenomic data-driven and well supported domain chronology [21].

## Materials and methods

Disorder analysis was conducted for two datasets corresponding to increasingly more granular levels of complexity in a biological system, namely, proteomes and protein domains. Our objective at the organismal level was two-fold: (i) compute disorder in a set of representative proteomes, and (ii) analyze protein domains using established protocols and construction of an evolutionary “reference” based on a proteomic consensus [15, 16]. The proteome data was derived from a phylogenomic reconstruction of an evolutionary timeline of domains defined at fold family level of SCOP classification [21]. Leveraging hidden Markov Models in the SUPERFAMILY database [32, 33], we analyzed domain family assignments (e-value < 0.0001) in the proteomes of 8,127 completely sequenced cellular and viral genomes. Domains were labeled with SCOP *concise classification strings* (*ccs*). Cellular proteomes comprised 139 archaeal, 1,734 bacterial, and 210 eukaryal representative and reference proteomes, sampling organisms in all major taxonomic groups of the RefSeq database [34] (Table 1). Due to the small size of viral proteomes, we selected all 6,540 nonredundant proteomes present in the NCBI viral genomes project (as of January 2019), out of which, HMMER assigned domain family occurrence and abundance in 6,044 proteomes [35]. The viral proteomes were further categorized using the Baltimore classification [36], which clusters viruses into groups on the basis of genome types and viral replication strategies. The domain *use* (number of unique domain families in a proteome) and *reuse* (total number of domains in families) values for each proteome were then computed. In addition to times of origin (*nd* values), a distribution of spread for domains in proteomes was computed via a fractional (*f*) value, which ranges from 0 (absent in all proteomes of the dataset) to 1 (present in all proteomes of the dataset).

**Table 1.** Proteomes downloaded from RefSeq [34] and the NCBI Viral Genomes Projects [34] presented by taxonomic group for phylogenomic and disorder analysis.

| Taxonomic Group | Reference | Representative | Group total |
| --- | --- | --- | --- |
| Archaea | 0 | 139 | 139 |
| Bacteria | 110 | 1,624 | 1,734 |
| Fungi | 1 | 47 | 48 |
| Invertebrates | 2 | 17 | 19 |
| Plants | 1 | 55 | 56 |
| Protozoa | 0 | 50 | 30 |
| Vertebrates | 3 | 54 | 57 |
| Viruses | 6,044* |  | 6,044 |
| Total |  |  | 8,127 |

\*Proteomes with HMMER assignments. The starting number of viral proteomes was 6,540. This included all complete RefSeq proteomes in the NCBI Viral Genomes Project (January 2019), not only limited to representative or reference categories, but excluding delta viruses and unclassified viruses.

A total of 103,000 protein sequences available out of the 110,800 structures corresponding to SCOP domains were downloaded from the ASTRAL compendium (version 1.75). The sequences were annotated with SCOP domain families [37] along with their times of origin from a chronology of domains constructed from the 8,127 proteomes [21] that were indexed with molecular function information [38] catalogued into 7 broad and 49 detailed function categories. Structural disorder was calculated using a standalone downloadable version of IUPRED, using the ‘long’ disorder option [39]. A residue was considered locally disordered if it scored above the cutoff value of 0.5 and disorder of the protein was computed as the ratio of the constituent disordered residues. The mean disorder for each proteome was calculated as the mean of disordered fraction percentage of proteins [40]:

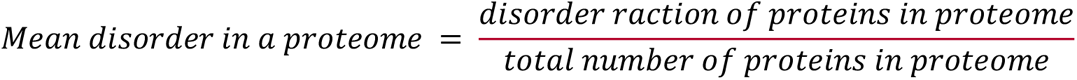

Similarly, the mean disorder for each SCOP domain family was calculated as the average of the disorder percentage of corresponding ASTRAL domain sequences:

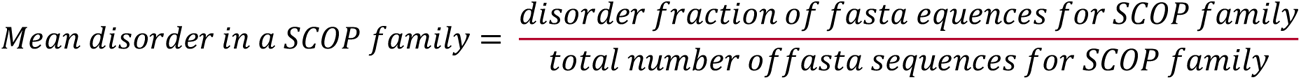

A proteome or domain family was classified as belonging to the ‘ordered’, ‘moderate disorder’, or ‘high disorder’ categories, if their mean disorder score ranged between 0–0.1, 0.1– 0.3, or greater than 0.3, respectively [41].

We provide graphical summaries of data distributions with boxplots, violin plots and boxenplots. We note that boxenplots, also known as letter-value plots [42], are useful for representing large datasets that are not normally distributed and have fat-tailed distributions.

## Results

### Intrinsic disorder of structural domains analyzed at proteome level

We surveyed intrinsic disorder in 8,127 proteomes belonging to viruses and organisms representing the three cellular superkingdoms (Archaea, Bacteria and Eukarya). We calculated a disorder score as a fraction of disordered residues in structural domains defined at fold family level of SCOP classification [37] and at proteome level. We refer to this fraction as ‘mean disorder’ throughout the manuscript. In cellular organisms, mean disorder showed a gradual increase with mean protein length and total number of proteins present in proteomes (Fig. 1). In contrast, mean disorder of viral proteomes decreased with increases in both of these parameters. There was a significant difference among the means of the disorder distribution among superkingdoms and viruses, indicating each broad taxonomic group had its own profile of disorder (Fig. 2a). Mean disorder decreased with increased use and reuse of domains in viral proteomes, while the opposite was observed for cellular proteomes, leading to tight clusters of proteomes into their respective taxonomic groups (Fig. 2b). Proteomes with mean disorder scores corresponding to the ‘ordered’ category demonstrated a high domain use and reuse (Fig. 2c). While proteomes with moderate disorder were distributed with a lower median domain use and reuse, the distribution was positively skewed with scores comparable to those of ‘ordered’ proteomes.

**Figure 1.**
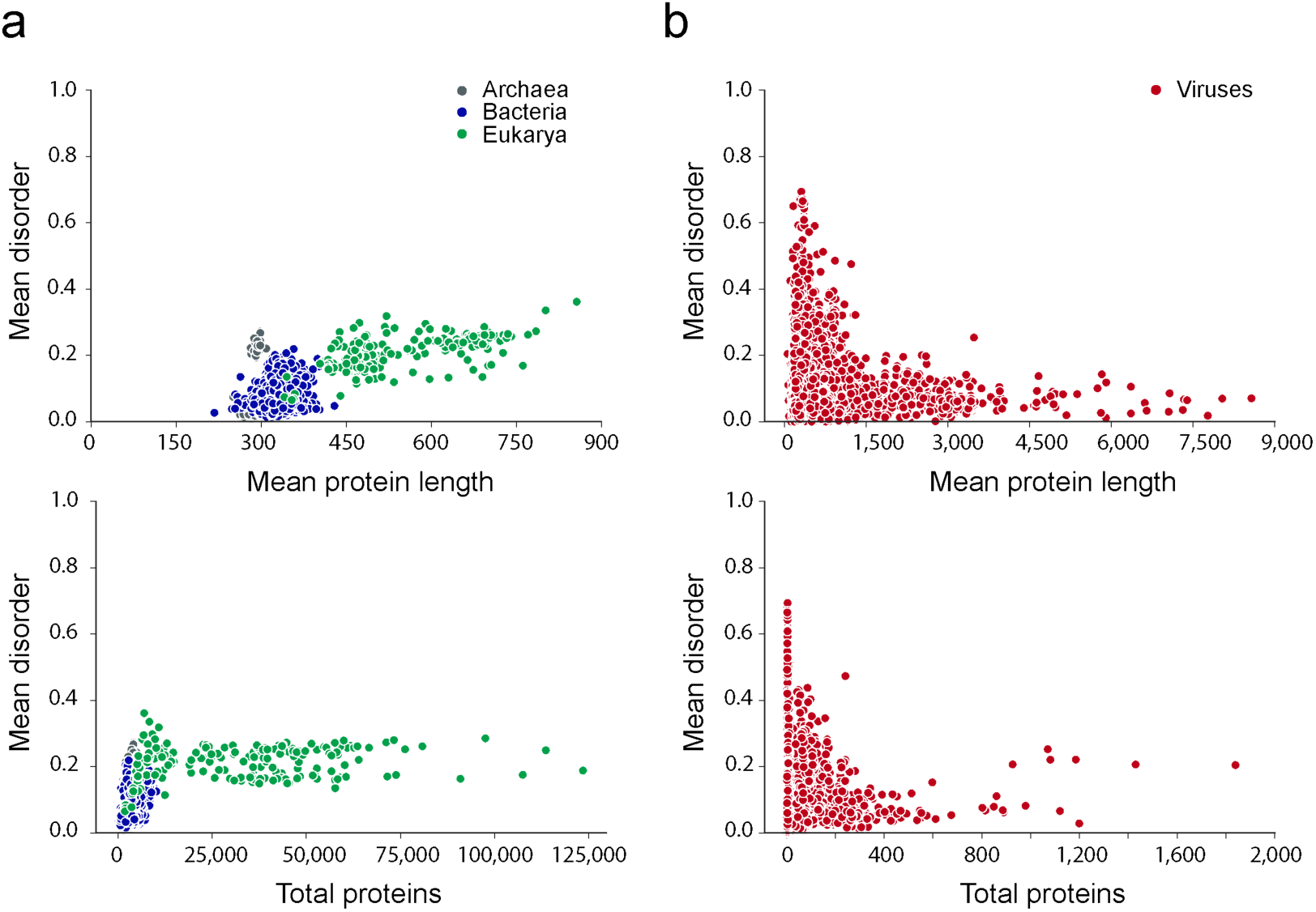
Relationship of disorder with protein length and proteomic complexity in organisms of superkingdoms and viruses. (a) Mean disorder by mean protein length and total number of proteins in the sampled cellular proteomes. (b) Mean disorder by mean protein length in the sampled viral proteomes.

**Figure 2.**
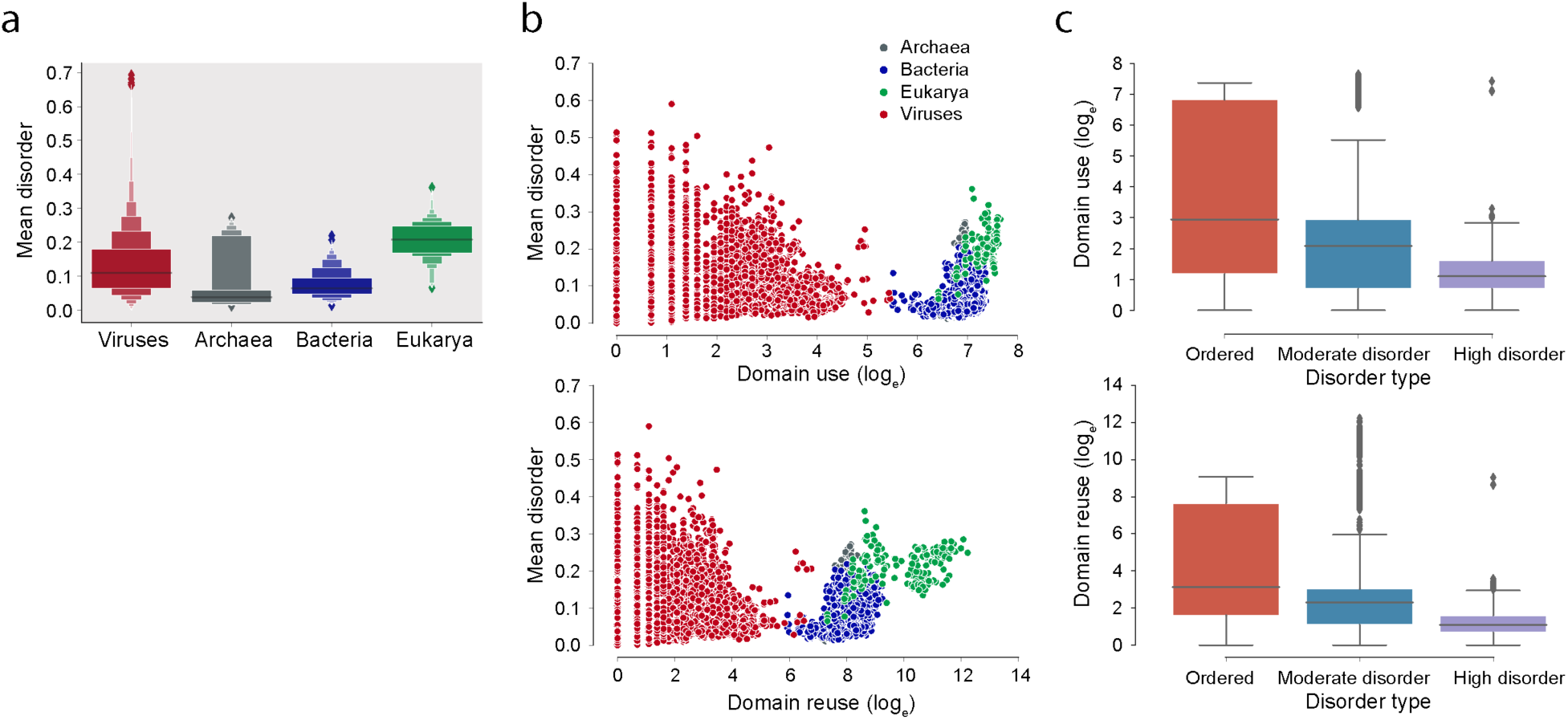
Disorder in cellular and viral proteomes. (a) Boxenplots show disorder scores by superkingdom (Kruskal-Wallis H-test corrected p-value = 1.162e-224 using the Holm-Bonferroni method at Family-wise Error Rate = 0.05). (b) Scatterplots showing the relationship between mean disorder and domain use and reuse in the three superkingdoms of life and viruses. (c) Distribution of domain use and reuse (logarithmic scale) in proteomes grouped by mean disorder categories.

Most archaeal proteomes fell in the category of ‘ordered’ proteomes, all forming overlapping clusters in scatterplots, with the exception of those in class *Halobacteria,* a member of the archaeal phylum *Euryarchaeota* (Fig. 3). The halobacterial proteomes exhibited moderate mean disorder scores, high domain use and reuse, and relatively larger proteome sizes. Among bacterial proteomes, all superphyla had proteomes that possessed ‘moderate disorder’ (Fig. 4a), with the bulk of them having proteomes larger that archaeal proteomes (Fig. 4b) with low disorder levels belonging to the ‘ordered’ category (Fig. 4c). Unlike archaeal proteomes, bacterial proteomes had no distinct clustering based on both mean disorder and domain use and reuse (Fig. 4d). In sharp contrast, most eukaryotic proteomes had moderate disorder levels (Fig. 5a), with the exception of a few fungi and protozoan species holding proteomes with ordered and high disorder levels (Table 2). Eukaryotic proteomes clustered with their respective taxonomic groups based on domain use and formed overlapping clusters based on domain reuse (Fig. 5b).

**Figure 3.**
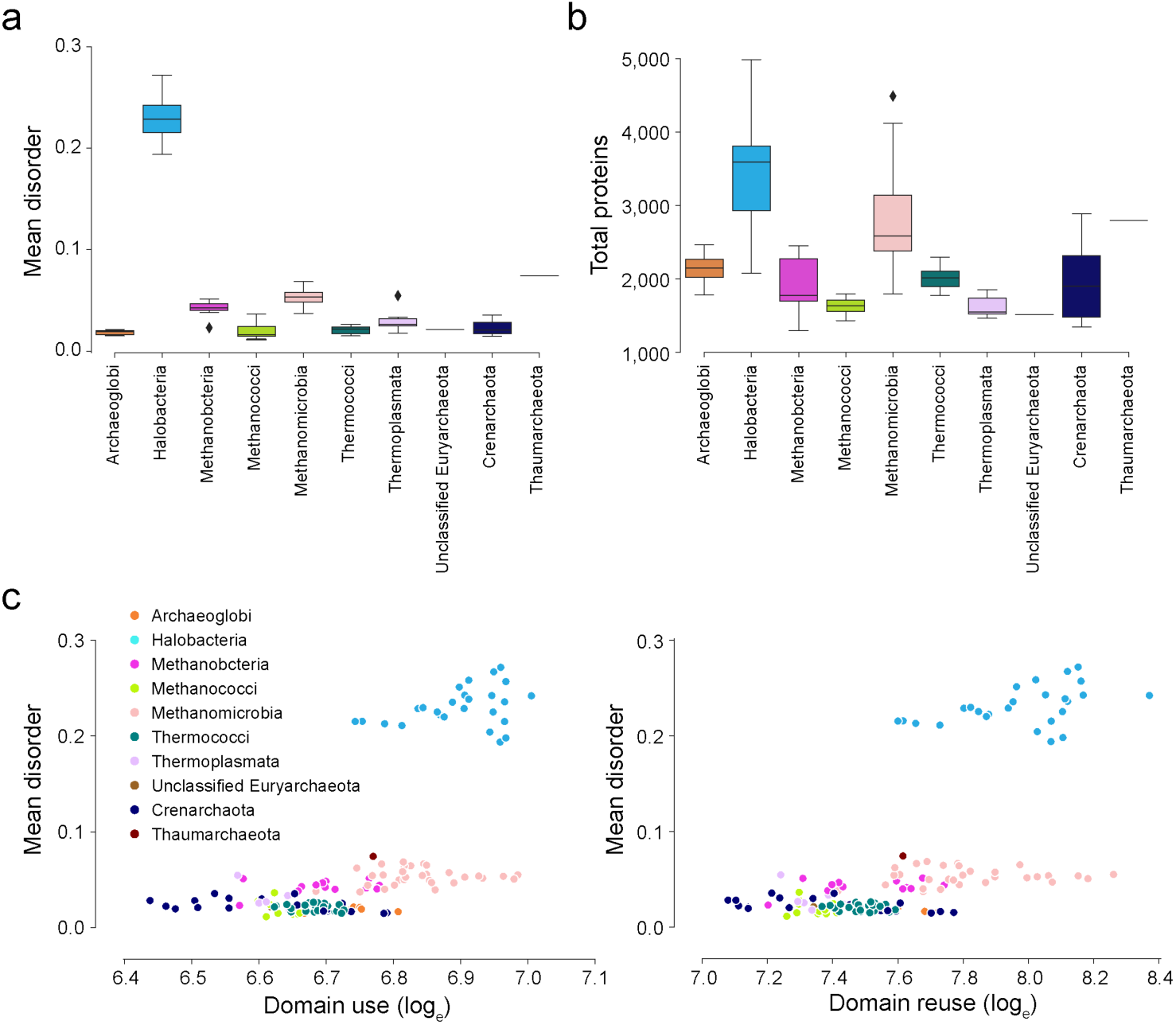
Disorder in archaeal proteomes. (a) Distribution of mean disorder in sampled archaeal proteomes divided into taxonomic classes. (b) Distribution of total proteins in sampled archaeal proteomes grouped by classes. (c) Scatterplot between mean disorder and domain use and reuse (logarithmic scale) in sampled archaeal proteomes grouped by classes.

**Figure 4.**
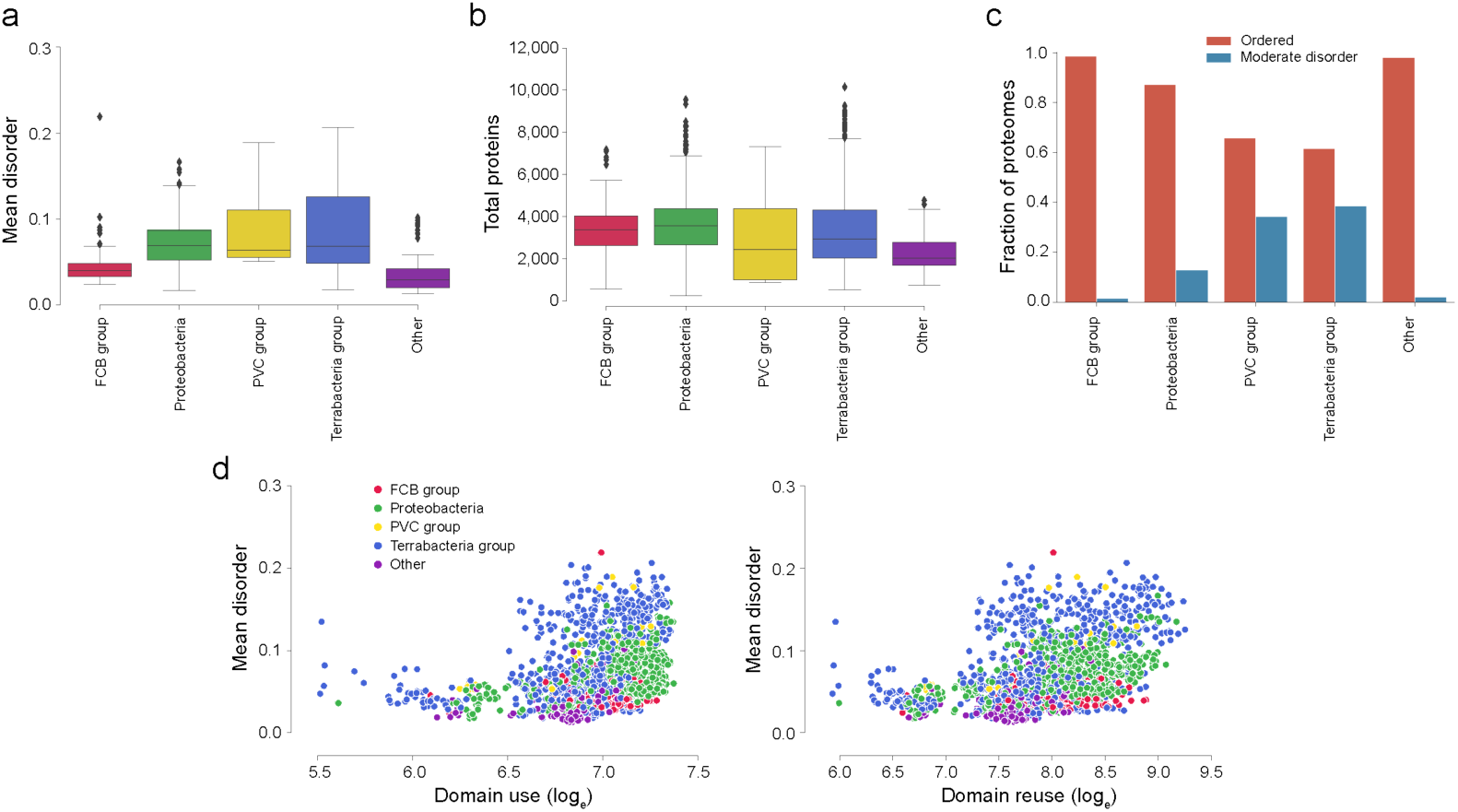
Disorder in bacterial proteomes. (a) Distribution of mean disorder in sampled bacterial proteomes grouped by superphyla. (b) Distribution of total proteins in sampled bacterial proteomes grouped by superphyla. (c) Distribution of bacterial proteomes in disorder groups by superphyla. (d) Scatterplots between mean disorder and domain use and reuse (logarithmic scale) in sampled bacterial proteomes grouped by superphyla.

**Figure 5.**
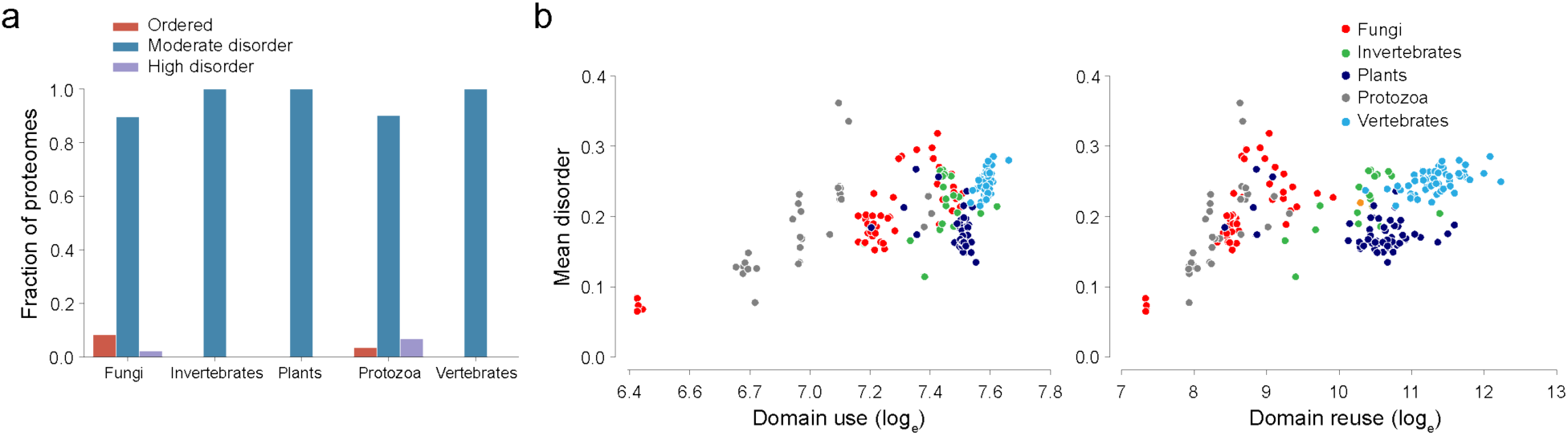
Disorder in eukaryotic proteomes. (a) Distribution of sampled eukaryotic proteomes into disorder groups by major taxonomic groups. (b) Scatterplot between mean disorder and domain use (logarithmic scale) in sampled eukaryotic proteomes grouped by major taxonomic groups. (c) Scatterplots between mean disorder and domain use and reuse (logarithmic scale) in sampled eukaryotic proteomes grouped by major taxonomic groups.

**Table 2.** Eukaryotic proteomes in the ordered and high disorder categories.

| Organism | Taxonomic group | Total proteins | Mean protein length | Mean disorder | Disorder category |
| --- | --- | --- | --- | --- | --- |
| *Encephalitozoon cuniculi* | Fungi | 1996 | 359.1 | 0.084 | ordered |
| *Encephalitozoon hellem* | Fungi | 1847 | 357.0 | 0.068 | ordered |
| *Encephalitozoon intestinalis* | Fungi | 1938 | 341.0 | 0.074 | ordered |
| *Encephalitozoon romaleae* | Fungi | 1831 | 353.9 | 0.065 | ordered |
| *Neurospora crassa* | Fungi | 10812 | 520.9 | 0.318 | high disorder |
| *Babesia microti* | Protozoa | 3601 | 439.8 | 0.078 | ordered |
| *Neospora caninum* | Protozoa | 6934 | 856.4 | 0.361 | high disorder |
| *Toxoplasma gondii* | Protozoa | 8318 | 801.6 | 0.335 | high disorder |

Viral proteomes constituted the majority of highly disordered proteomes present in our dataset. They exhibited high levels of mean disorder, with the proteomes of ssDNA viruses being the most disordered (Fig. 6a). The fraction of moderately disordered proteomes, when compared to the other two categories, was greater for viruses in each viral replicon group, with the exception of dsRNA and plus-ssRNA viruses, which had a higher number of ordered proteomes (Fig. 6b). Remarkably, viruses and in particular ssRNA-RT viruses had the largest fraction of high disorder proteomes in all cellular and viral groups. An inspection of viruses grouped according to the superkingdom of their hosts revealed interesting patterns. Archaeoviruses had a high number of ordered proteomes followed by high disorder proteomes (Fig. 6b). On the other hand, high disorder proteomes were absent in bacterioviruses, with the majority of the proteomes possessing moderate disorder. Similarly, eukaryoviruses had a similar fraction of proteomes with moderate disorder but exhibited a small fraction of proteomes with high disorder. Among the eukaryoviruses with high disorder, eukaryoviruses infecting invertebrate and plant (IP) hosts represented the highest fraction followed by eukaryoviruses with metazoan (vertebrates, invertebrates, and humans), plant (all plants, blue-green algae, and diatoms) and fungal hosts (Fig. 6b). As noted earlier, a tendency of decreasing spread of disorder scores with increasing domain use and reuse and proteomic sizes manifested for viruses belonging to all replicon groups (Fig. 6c). In particular, the bulky proteomes of dsDNA viruses showed a decrease in disorder scores with increasing domain use and reuse, while smaller viral proteomes belonging to other replicons exhibited widespread disorder.

**Figure 6.**
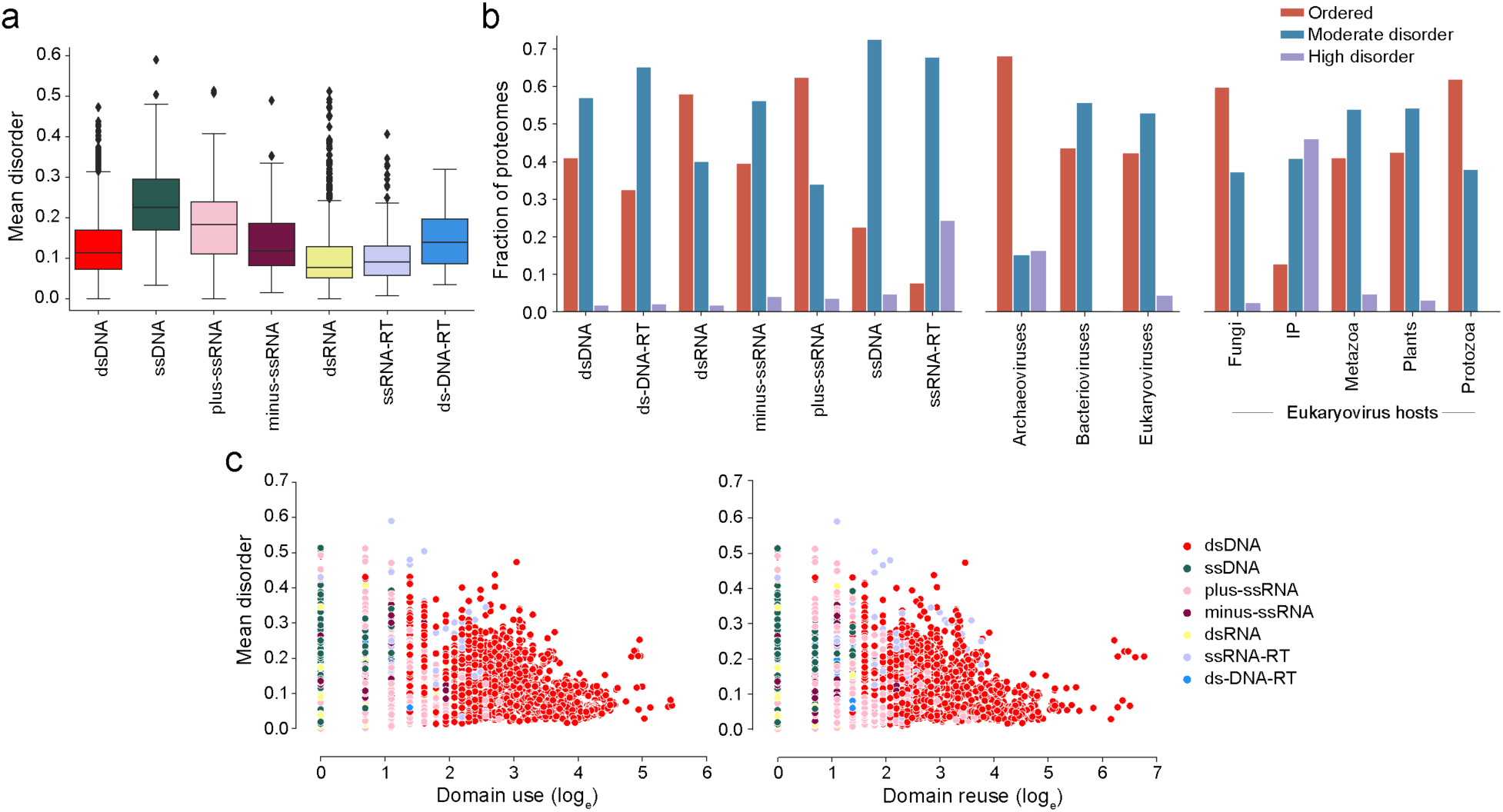
Disorder in viral proteomes. (a) Distribution of mean disorder in sampled viral proteomes grouped by replicon type according to the Baltimore classification of viruses [35]. (b) Distribution of sampled viral proteomes based on replicon type, host superkingdoms, and eukaryovirus hosts. Eukaryovirus hosts were classified into Protista (animal-like protists), Fungi, Plants (all plants, blue-green algae, and diatoms), Invertebrates and Plants (IP), and Metazoa (vertebrates, invertebrates, and humans). Host information was available for 5,959 of the 6,044 viruses sampled in this study. (c) Scatterplots between mean disorder and domain use and reuse (logarithmic scale) in sampled viral proteomes grouped by replicon type.

### Evolution of intrinsic disorder analyzed at structural domain level

To study intrinsic disorder evolving in the universe of structural domains, we traced the number of domains exhibiting order, moderate disorder, and high disorder levels along an evolutionary chronology of first appearance of domains in proteomes [21]. This chronology has been benchmarked by about two decades of research [15, 18, 20, 21, 30, 43–45]. The time of origin of structural domains was derived directly from a phylogenomic tree of domains defined at SCOP fold family level of abstraction [21]. Time of origin was expressed as a *node distance* (*nd*), with *nd* = 0 corresponding to the origin of protein domains and *nd* = 1 corresponding to the present. In some plots, time of origin was binned in 0.1 intervals. Fig. 7 describes the accumulation of ordered, moderate disorder, and high disorder domains in evolution. A comparative genomic analysis of domains in the proteomes of organisms in superkingdoms and viruses revealed similar distribution patterns of mean disorder (Fig. 7a). However, ordered domains appeared early in evolution, followed by domains with moderate disorder and high disorder, in that order (Fig. 7b). The early origin of ordered domains is also demonstrated by the timeline of domains shared among cells and those shared between cells and viruses on the basis of disorder category (Fig. 7c). The bulk of the domains with high disorder for both sets appeared later in the timeline and followed the same progression of times of origin observed at global level (Fig. 7b). The evolutionary accumulation of domains followed a pattern similar to that of their origin (Fig. 7d). Ordered domains appeared first and accumulated the most along the timeline, followed by domains with moderate disorder and finally domains with high disorder. In fact, there was a ‘big bang’ of innovation associated with ordered domains that manifested with a significant accumulation peak appearing halfway in the evolutionary timeline (Fig. 7d). In contrast, the evolutionary growth of domains exhibiting moderate and high disorder was limited, with the late-appearing highly disordered domains showing the slowest pattern of evolutionary growth. The spread of mean disorder gradually increased in evolution, first in domains with moderate disorder and then in high disorder domains and was more pronounced as time progressed, especially in domains with high disorder (Fig. 7e). We also found there was a gradual decrease in mean disorder with increases in average domain length (Fig. 7f), with domain shared only among cells and those shared between viruses and cells demonstrating a similar behavior. This clear association between mean disorder and polymer length was paraphrased with that found between mean disorder and mean protein length in viruses (Fig. 1b), showing that the culprit manifests in domains, linker and terminal regions of the proteins.

**Figure 7.**
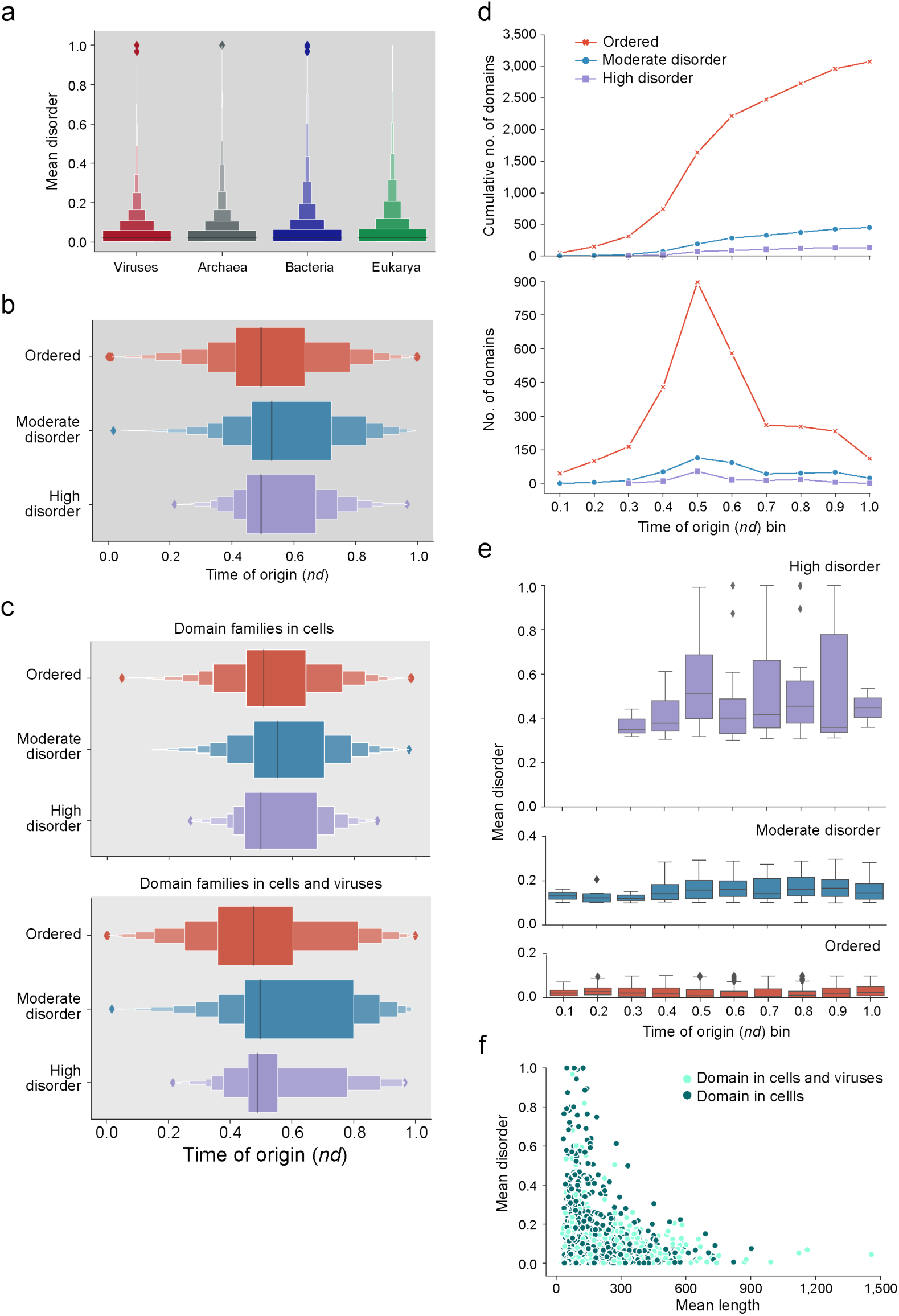
Evolution of intrinsic disorder in the structural domains of proteins. (a) Mean disorder in domains present in superkingdoms and viruses. (b) Evolutionary history of domains corresponding to the three disorder groups. (c) Evolutionary history of domains in cells and in cells and viruses. (d) Number of domains expressed in absolute and cumulative numbers that appear in the bins of the evolutionary timeline. (e) Distribution of disorder along the evolutionary chronology of domains grouped by types of disorder. Note that families with high disorder appear in the third time of origin (*nd*) bin of the chronology. (f) Relationship between mean disorder and mean length of domains shared only among cells and those shared between viruses and cells.

An inspection of evolutionary history of domains indexed with disorder categories and belonging to Venn groups describing the sharing of domains across supergroups Archaea, Bacteria, Eukarya and viruses revealed that ordered domains were the first to appear in all 15 Venn taxonomic groups (Fig. 8). Note that the appearance of Venn groups along the evolutionary timeline occurred in a specific order delimiting six evolutionary phases and two universal ancestors of life, the last universal common ancestor (LUCA) and the last universal cellular ancestor (LUCellA) of life. Some implications of phases and ancestors were recently discussed [46]. Phase 0 (communal world) was only populated by universal domains common to all supergroups (the ABEV group). The phase included the most ancient domain family, the ‘ABC transporter ATPase domain-like’ (SCOP *ccs*: c.37.1.12), which belonged to the set of ordered domains, and “Cold shock DNA-binding domain-like” family (b.40.4.5), which was the most ancient among moderately disordered domains. The end of this phase defines the proteome of LUCA encompassing ordered domains and a minority of moderate disorder domains. Phase I (rise of viral ancestors) was populated by universal domains and by a minority of families common to all superkingdoms. The end of this phase defined the proteome of LUCellA. Phase II (birth of archaeal ancestors) comprised 4 Venn groups (ABEV, ABE, BEV, and BE) and defined an ancestral stem line of cellular descent, which ultimately gave rise to Archaea. The first high disorder domain, the ‘HSP40/DnaJ peptide-binding domain’ family (b.4.1.1) belonging to the ABEV group, appeared during this phase. Phase III (diversified Bacteria) harbored more than half of the Venn groups, all of them involving Bacteria. Finally, Phases IV and V introduced the rest of the Venn groups, including virus-specific domains (Venn group V), which appeared at the beginning of Phase IV. The distribution of domains indexed with molecular functions along the evolutionary timeline shows an early evolution of ordered domains in all seven major functional categories, except for ‘Information’, for which domains with moderate disorder evolved earlier than ordered domains (Fig. 9). The latest domains to be introduced in the timeline belonged to the category of ‘Extracellular processes’ and ‘Other’.

**Figure 8.**
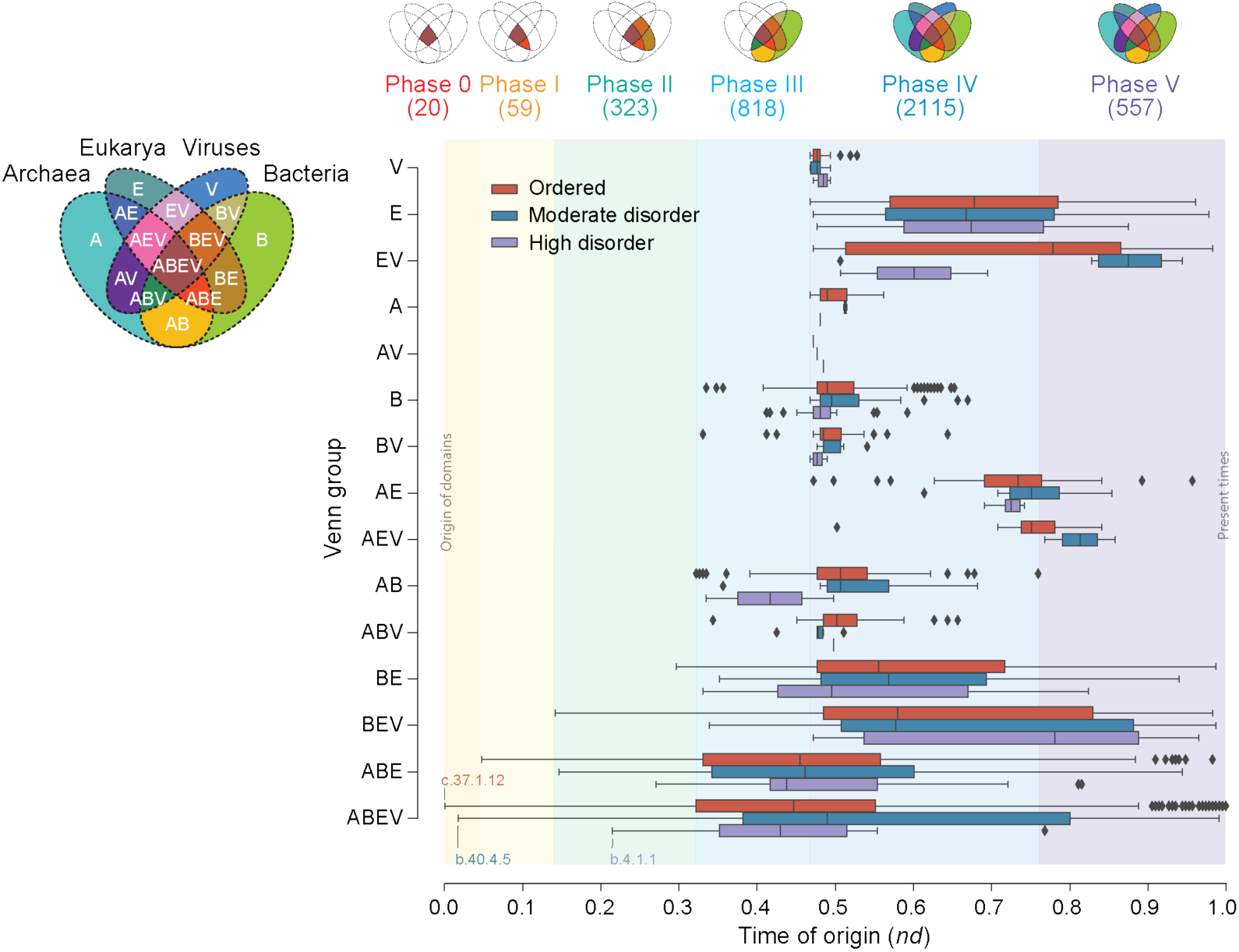
Phylogenomic analysis of domains in cells and viruses. Venn diagrams describe the distribution of domains among supergroups Archaea, Bacteria, Eukarya and viruses and the appearance of Venn taxonomic groups along a chronology of domains with the temporal dimension describing their time of origin (*nd*). Venn-group colors reflect the evolutionary chronology of Venn-group appearance, which defines 6 evolutionary phases. Domains appearing in each phase are given in parentheses. The evolutionary history of domains classed into disorder groups is depicted for each Venn distribution group along the chronology with box plots.

**Figure 9.**
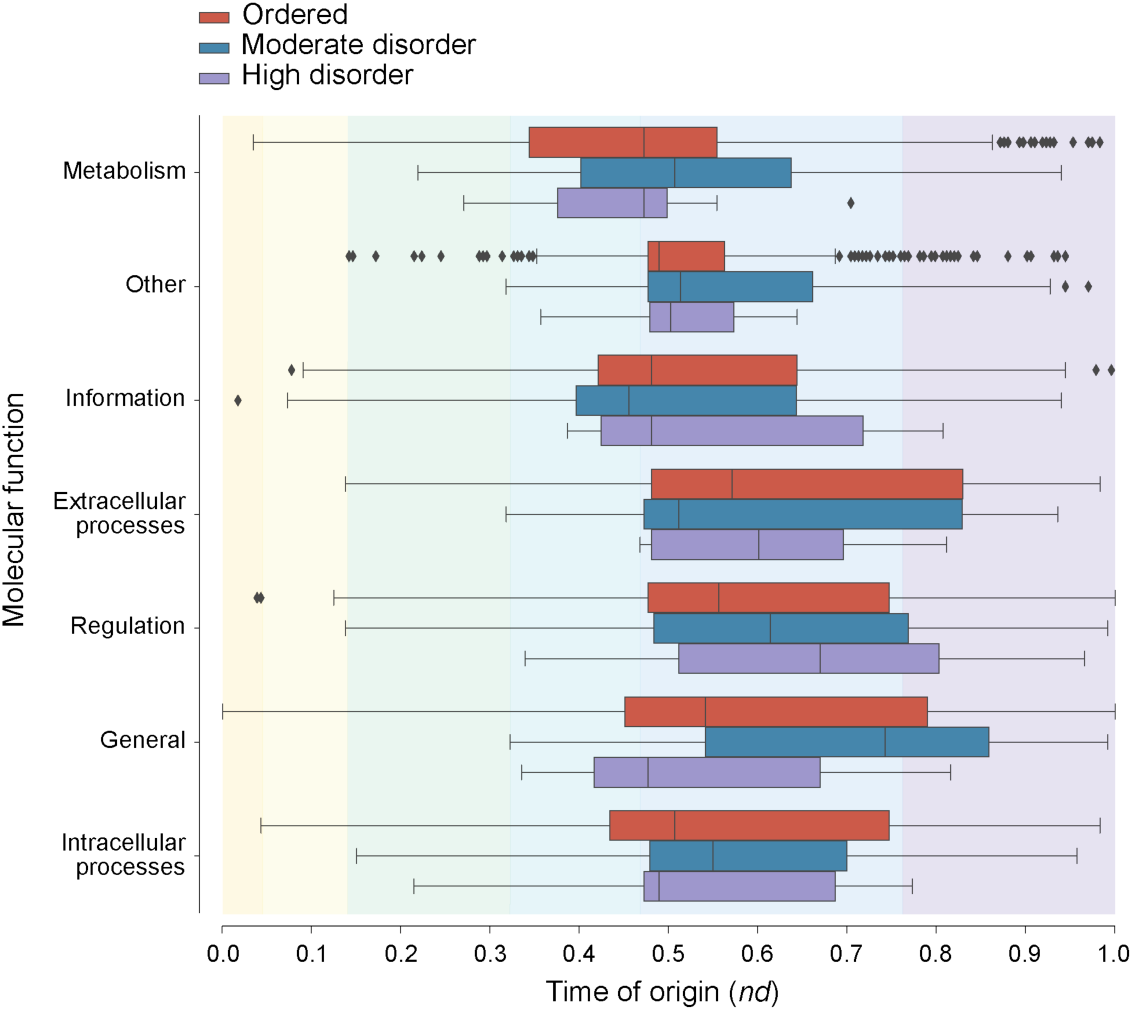
Evolutionary history of domains grouped by molecular function and disorder category.

The Venn diagrams *per se* dissected in a comparative proteomic exercise by the categories of disorder revealed domain sharing patterns across superkingdoms and viruses that were indicative of the origin of disorder (Fig. 10). While the highest number of ‘ordered’ (27.5%) and ‘moderate disorder’ (22.8%) domains belonged to the ABEV Venn group, only a tenth of all ‘high disorder’ (9.9%) domains belonged to that group and their number was not the highest in the set. The ABE and BE groups represented greater than one-third of the ‘high disorder’ domains (19.8% and 18.3%, respectively), suggesting the more derived origin of increased disorder levels. These findings align with the ancient origin of the ordered and moderate disordered groups directly inferred from the chronologies. We also mapped the spread of domains among proteomes (*f*-value) shared among cells and those shared between cells and viruses (Fig. 10) to investigate the predominant direction of gene transfer [19]. The number of ‘ordered’ domains shared with viruses were more widespread in proteomes than the domains shared only with cells (Fig. 10a), with significantly different means for each distribution. This points to the ancient origins of viruses, with the possibility of early viruses interacting with the last common ancestor of all superkingdoms [20]. Although ‘moderate disorder’ and ‘high disorder’ domains shared among cells appeared more widespread than those shared between cells and viruses, the means for either distribution in both sets were not significantly different (Fig. 10b and c). An inspection of Gene Ontology (GO) biological processes of ‘high disorder’ domains revealed that they were enriched in functions involving ‘Information’ mechanisms such as transcription and translation, in addition to domains involved in protein folding, photosynthesis, pathogenesis and viral processes (Tables 3 and 4). The widespread distribution of ordered domains in cells and viruses, along with the finding that viral proteomes are enriched in highly disordered domains, points to the transfer of genes from cells to viruses and/or that viruses evolved disorder independently as an optimization for fitness to compete with their hosts. We address these possibilities in the Discussion section. In particular, the spread of archaeal domains shared with viruses was significantly high in ‘ordered’ domains, especially domains involving ssDNA, dsRNA and plus-ssRNA viruses (Fig. 11). Most archaeal viruses belong to the dsDNA Baltimore groups, and a minority of them to the ssDNA group. They appear not to utilize RNA replicon strategies. Therefore, the spread of domains induced by RNA viruses in Archaea likely arises during their evolution from ancient cells and not from horizontal gene transfer events [20].

**Figure 10.**
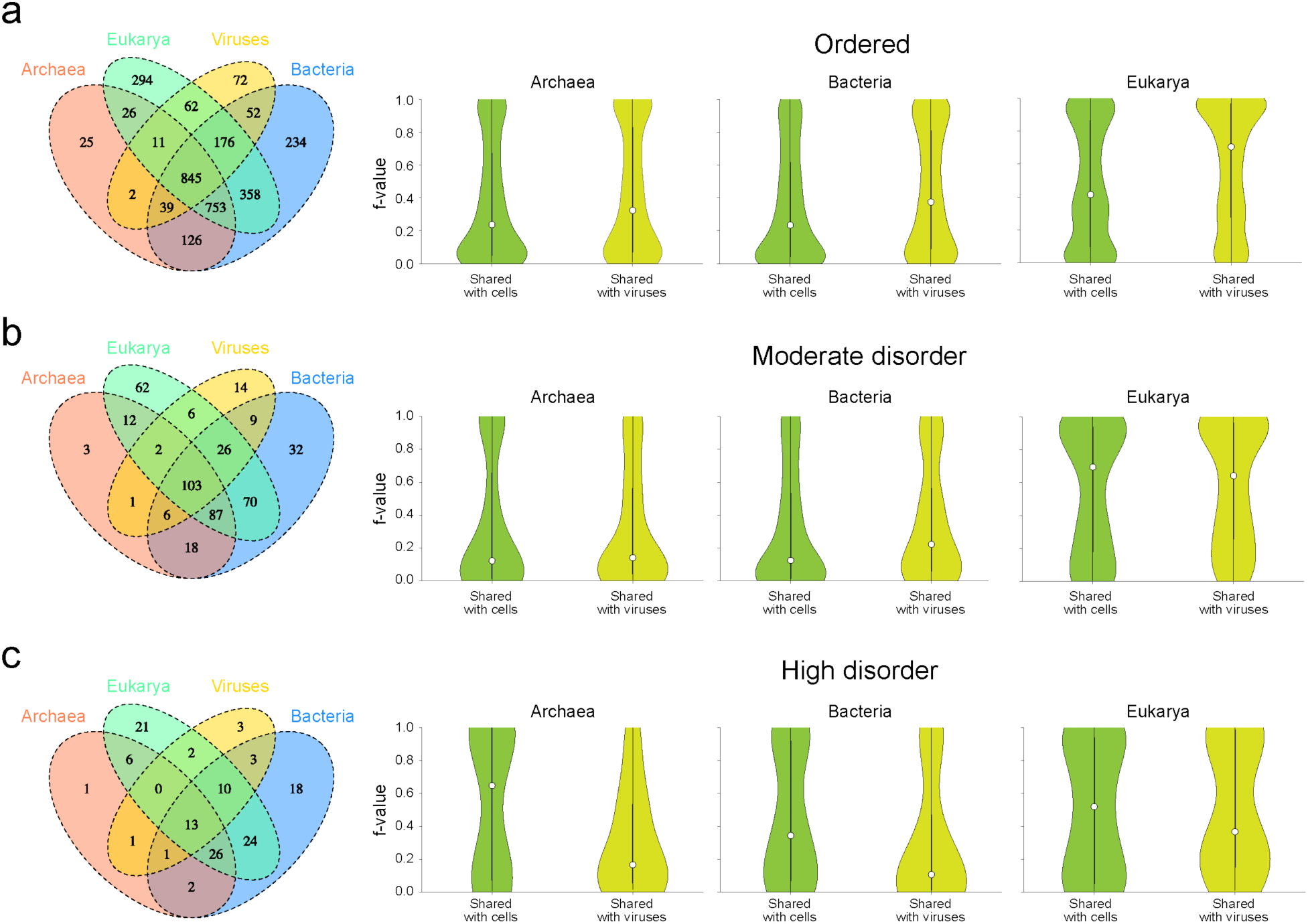
Sharing patterns and spread (*f*-value) of domains with indexed molecular functions in sampled proteomes classed by disorder group. Domains that were exclusive to each superkingdom and viral supergroup i.e. belonging to Venn groups A, B, E and V, were excluded from the spread distribution analysis. (a) Venn diagram and spread (*f-* value) of domains shared with cells and those shared with cells and viruses for ordered domains (Wilcoxon rank sum test, two-tailed, Archaea: *p*-value = 3.88 x 10^-5^; Bacteria: *p*-value = 2.12 x 10^-11^; Eukarya: *p*-value = 5.81×10^-17^). (b) Venn diagram and spread (*f-*value) of domains shared with cells and those shared with cells and viruses for moderately disordered domains (Wilcoxon rank sum test, two-tailed, fail to reject null hypothesis, Archaea: *p*-value = 0.756; Bacteria: *p*-value = 0.064; Eukarya: *p*-value = 0.251). (c) Venn diagram and spread (*f-*value) of domains shared with cells and those shared with cells and viruses for high disorder domains (Wilcoxon rank sum test, two-tailed, fail to reject null hypothesis, Archaea: *p*-value = 0.156; Bacteria: *p*-value = 0.0455; Eukarya: *p*-value = 0.567); *f*-value data from Mughal *et al.* [20].

**Figure 11.**
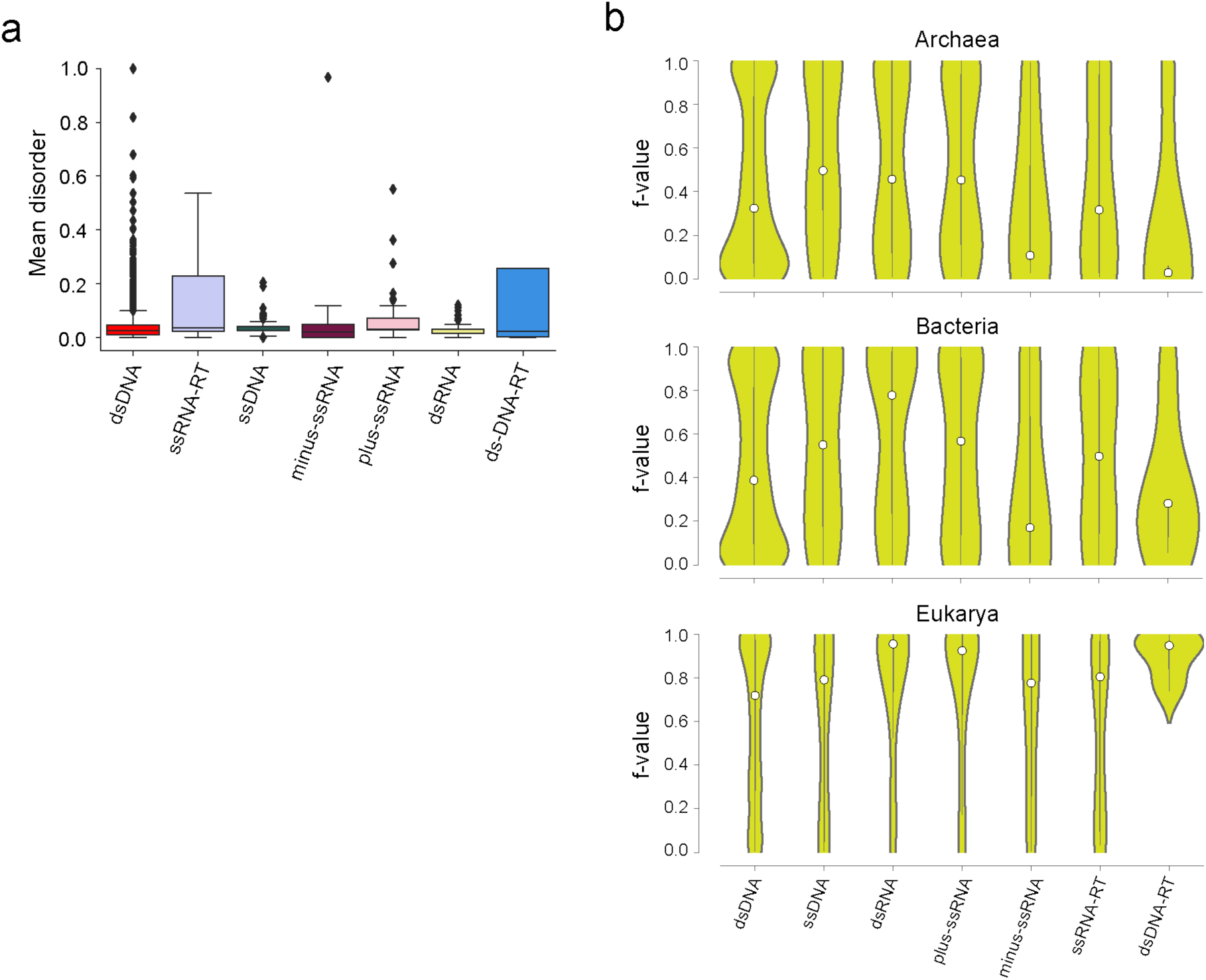
Disorder in viral domains. (a) Boxplots show mean disorder of 1,459 domains present in the sampled viral proteomes grouped by viral replicon type. (b) Violin plots show spread (*f*-value) of ordered domains that are shared between cells and viruses from Fig. 10 (*f*-value data from Mughal *et al.* [20]).

**Table 3.**
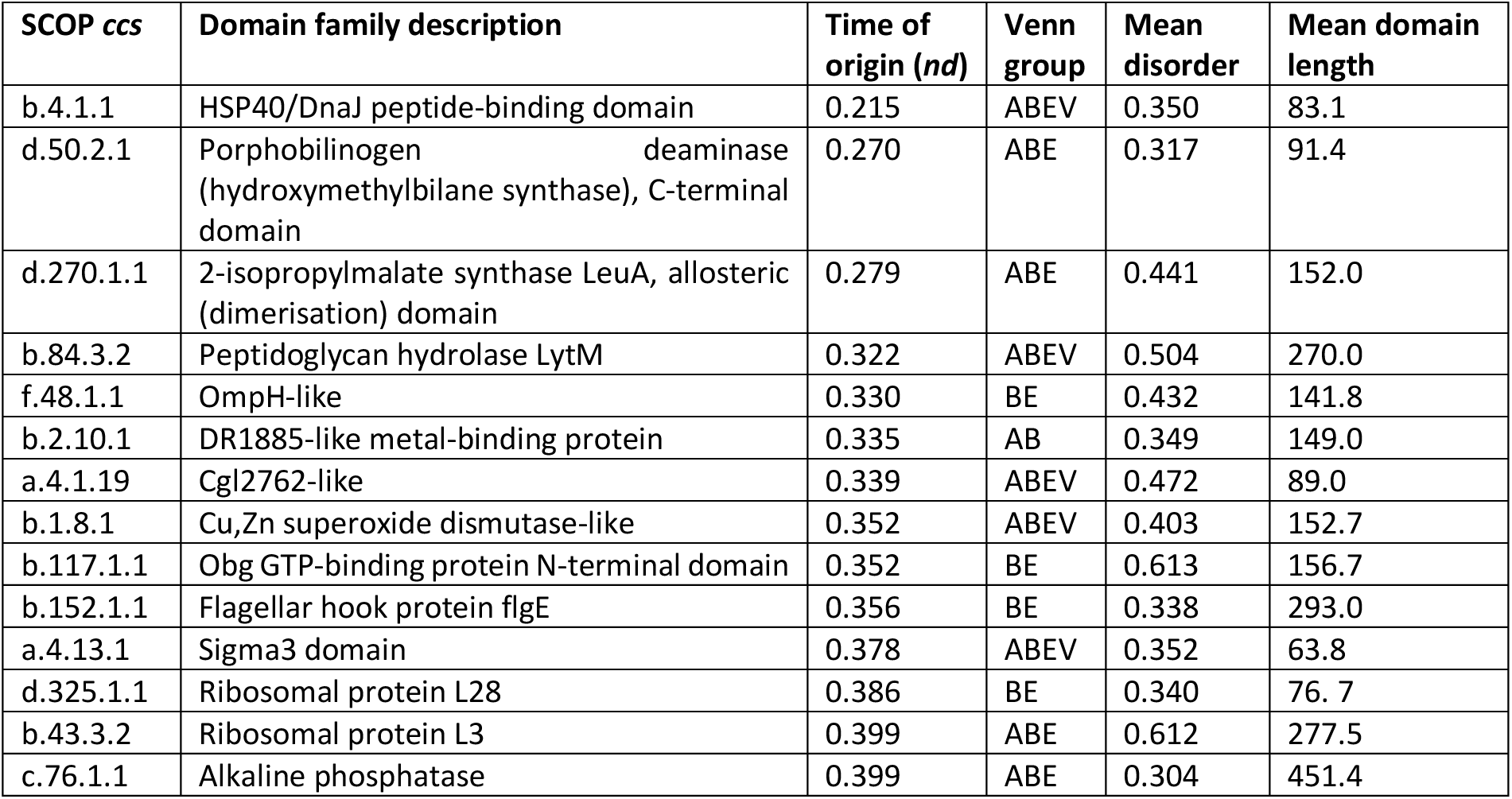
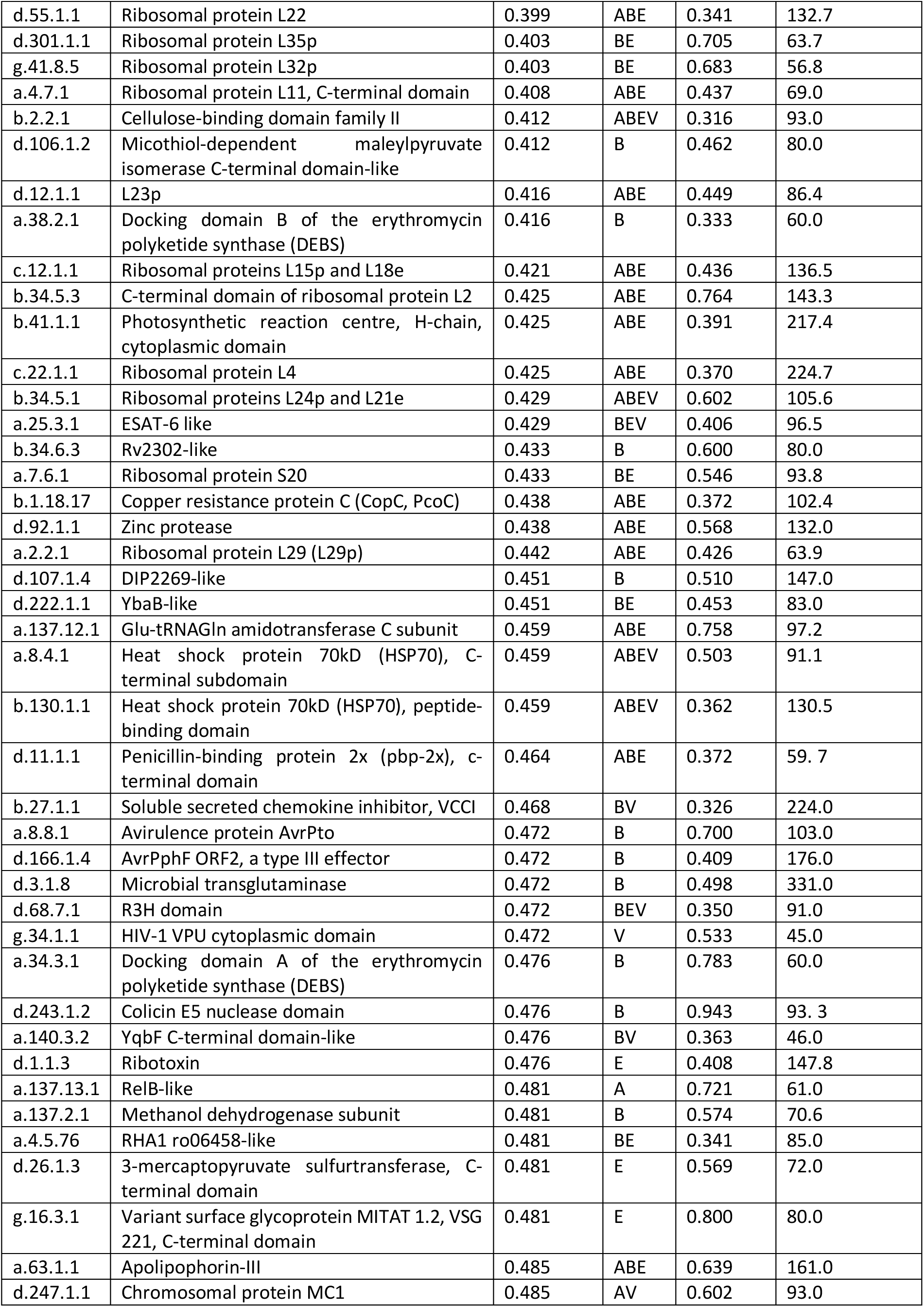
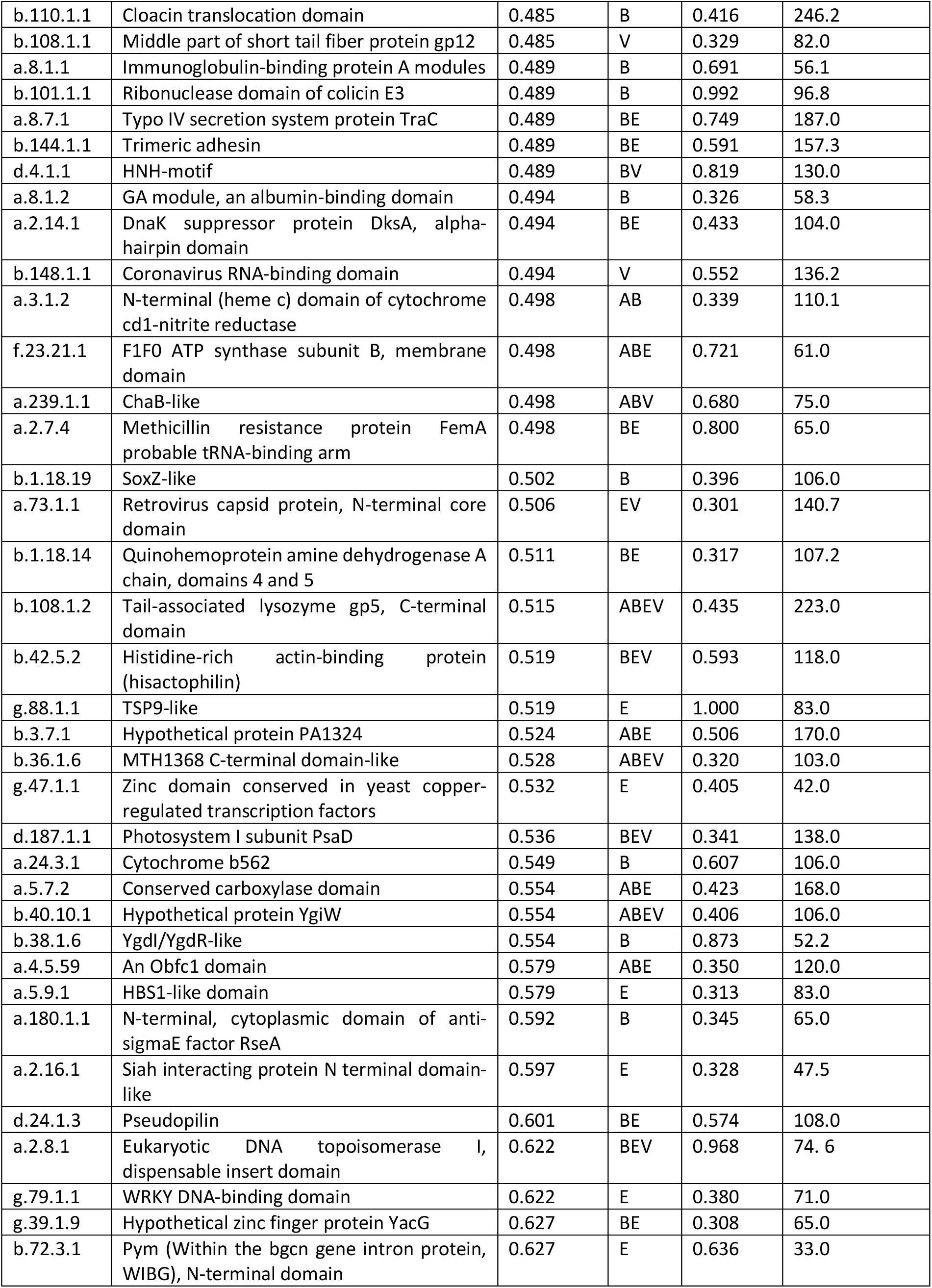
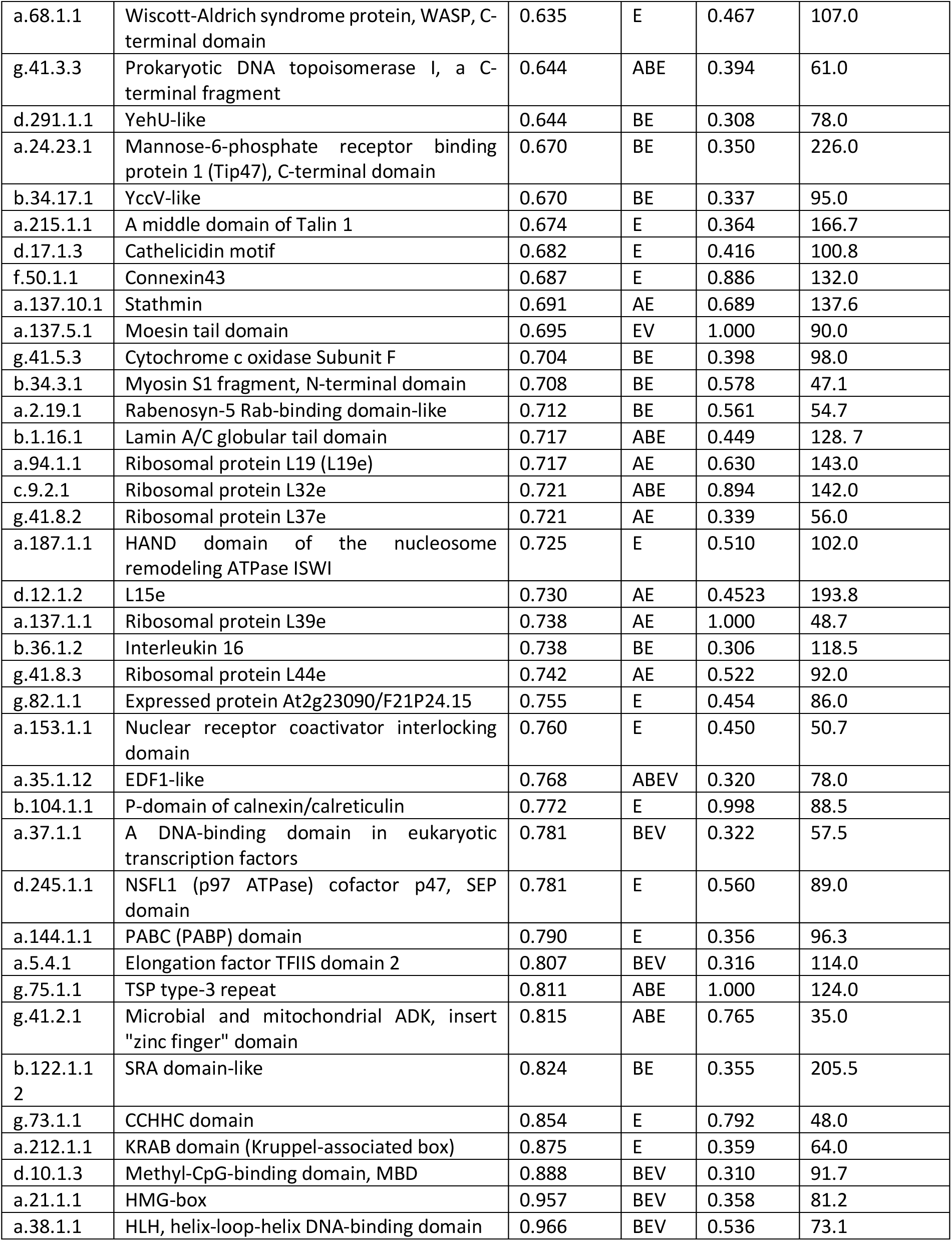
Structural domains with high mean disorder ordered according to their time of origin (*nd*).

**Table 4.**
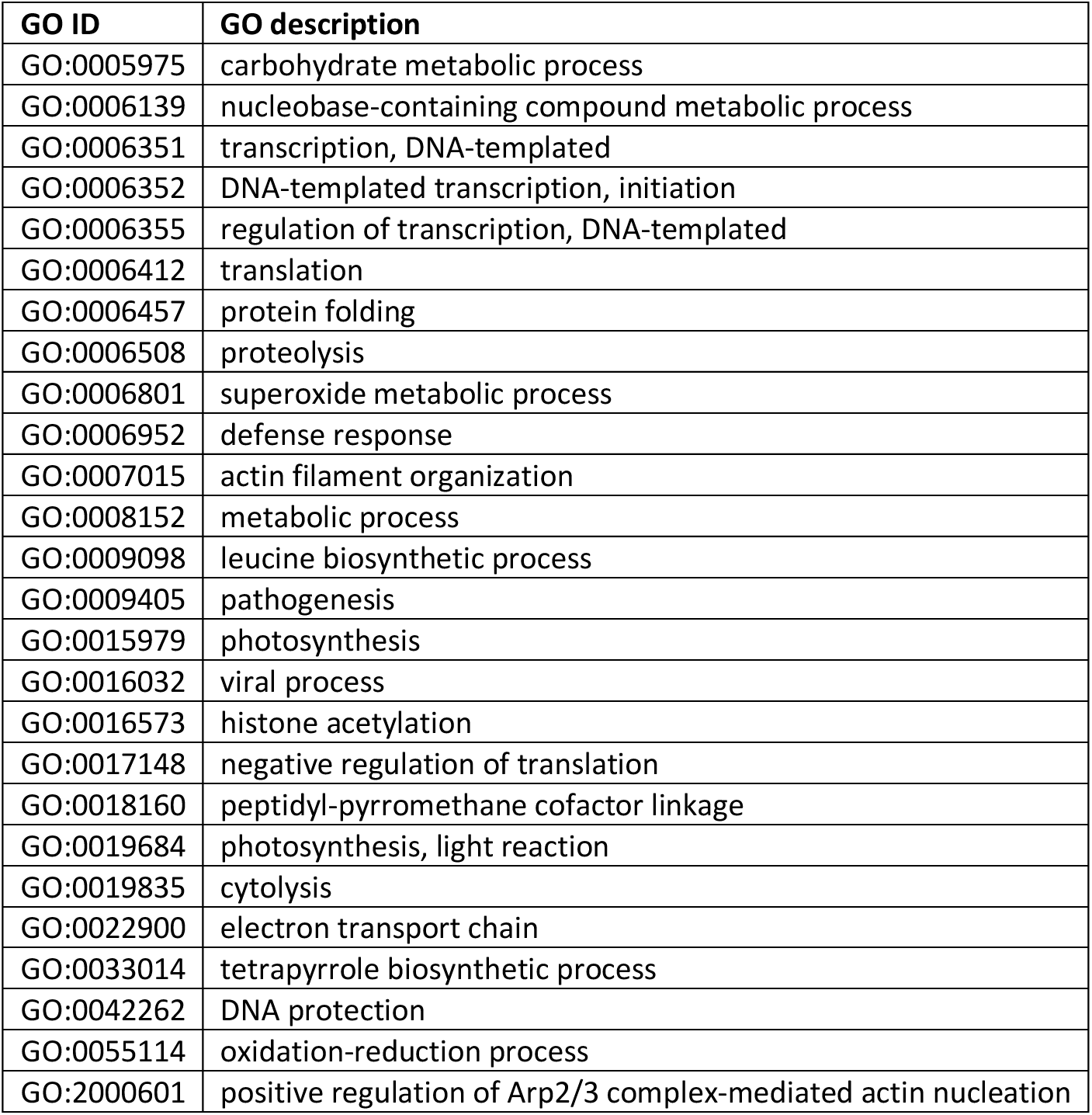
Highly enriched GO “biological process” terms in 131 domains with high disorder scores and a probability score equal to 1 in the Structural Domains Annotation Database (SDADB) [71].

| GO ID | GO description |
| --- | --- |
| GO:0005975 | carbohydrate metabolic process |
| GO:0006139 | nucleobase-containing compound metabolic process |
| GO:0006351 | transcription, DNA-templated |
| GO:0006352 | DNA-templated transcription, initiation |
| GO:0006355 | regulation of transcription, DNA-templated |
| GO:0006412 | translation |
| GO:0006457 | protein folding |
| GO:0006508 | proteolysis |
| GO:0006801 | superoxide metabolic process |
| GO:0006952 | defense response |
| GO:0007015 | actin filament organization |
| GO:0008152 | metabolic process |
| GO:0009098 | leucine biosynthetic process |
| GO:0009405 | pathogenesis |
| GO:0015979 | photosynthesis |
| GO:0016032 | viral process |
| GO:0016573 | histone acetylation |
| GO:0017148 | negative regulation of translation |
| GO:0018160 | peptidyl-pyrromethane cofactor linkage |
| GO:0019684 | photosynthesis, light reaction |
| GO:0019835 | cytolysis |
| GO:0022900 | electron transport chain |
| GO:0033014 | tetrapyrrole biosynthetic process |
| GO:0042262 | DNA protection |
| GO:0055114 | oxidation-reduction process |
| GO:2000601 | positive regulation of Arp2/3 complex-mediated actin nucleation |

To assess the relationship between mean disorder levels in viruses grouped according to their hosts, we examined the distribution of domains among the proteomes of archaeoviruses (*a*), bacterioviruses (*b*), and eukaryoviruses (*e*), which distribute into seven Venn taxonomic groups (Fig. 12a). Domains specific to one of the three viral groups make up the *a*, *b*, and *e* Venn groups, those shared by two viral groups make up the *ab*, *be*, and *ae* groups, and those shared by all three groups of viruses make up the *abe* core group. The *abe* core possessed the most ancient domains (Fig. 12b) and as expected comprised of ‘ordered’ and ‘moderate disorder’ domains (Fig. 12c). The origin of the *be, e, b* and *ab* groups followed in that order, respectively, for the most part matching the historical patterns observed in Fig. 8 for global Venn distribution groups (Fig. 12b). In the global analysis, the time of origin and temporal distribution of the ABE Venn group was followed by those of the BE, AB, B, A, E and AE groups, in that order. In the host-centric analysis of viral domains, a similar progression unfolds despite marked skewed distributions. For example, sections out of the centerline median of boxenplots, which contain 50% of the data, matched the expected progression. As expected, the distribution of disorder in these groups was much wider (Fig. 12c). In fact, domains specific to eukaryoviruses (*e* group) appeared to sample disorder throughout the scale, suggesting late but wide recruitment of high disorder domains prompted perhaps by a need to infect a wide range of hosts. Remarkably, the *e* group encompassed a total of 29 of the 37 known capsid protein domains, which appeared clustered in the timeline half way in evolution (*nd* = 0.468-0.524) (Table 5). Of the remaining capsid protein domains, five were present in the *b* Venn group, followed by 2 and 1 in the *be* and *abe* groups, respectively. Note that capsid domains belonging to the Venn group V and host-centric Venn groups *e* and *b* categories appeared earlier than other categories (beginning at *nd* = 0.468), supporting the origin and evolutionary transfer of domain novelties from viruses to hosts.

**Figure 12.**
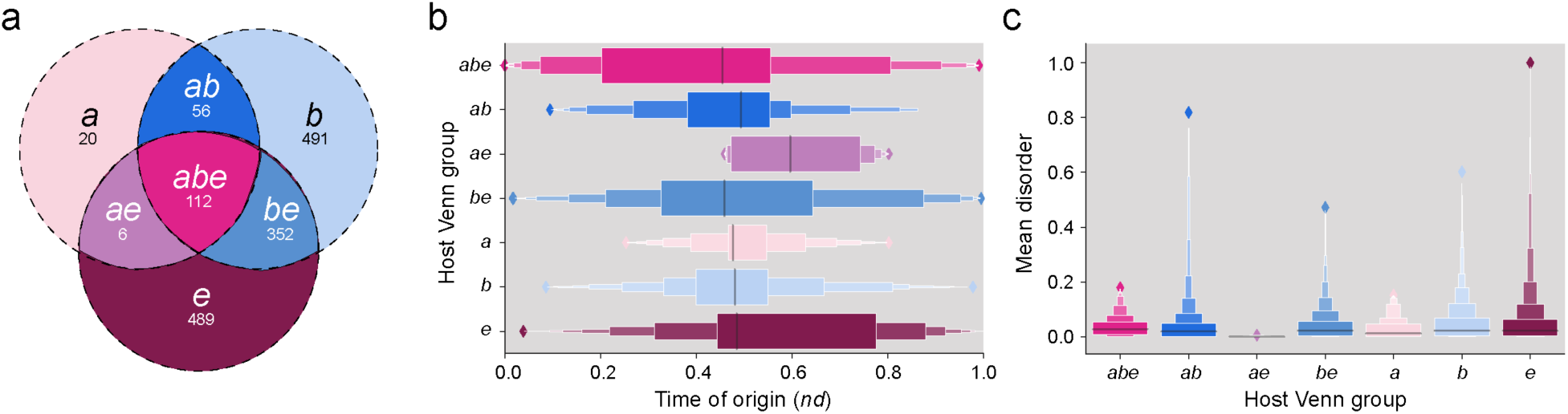
Domain disorder in the context of virus-host relationships. (a) Venn diagram showing the distribution of 1,526 domain families in archaeoviruses, bacterioviruses, and eukaryoviruses. Domains in the *abe* group do not imply that they were present in a virus that infects members of all three superkingdoms. The *abe* group only indicates the count of domains shared among archaeoviruses, bacterioviruses, and eukaryoviruses (domain counts data from Mughal *et al.* [20]). (b) Evolutionary history of domains in archaeoviruses, bacterioviruses, and eukaryoviruses. (c) Mean disorder distribution in archaeoviruses, bacterioviruses and eukaryoviruses. (c) Spread of mean disorder in domains of host Venn groups.

**Table 5.** Structural domains present in the proteins of viral capsids ordered according to their host-centric Venn group and time of origin (*nd*).

| SCOP ccs | Domain family description | Time of origin (*nd*) | Venn group | Host-centric Venn group |
| --- | --- | --- | --- | --- |
| d.183.1.1 | Major capsid protein gp5 | 0.519 | ABV | *abe* |
| d.85.1.1 | RNA bacteriophage capsid protein | 0.468 | V | *b* |
| a.84.1.1 | Scaffolding protein gpD of bacteriophage procapsid | 0.476 | V | *b* |
| h.1.4.1 | Inovirus (filamentous phage) major coat protein | 0.476 | BV | *b* |
| b.37.1.1 | N-terminal domains of the minor coat protein g3p | 0.476 | BV | *b* |
| b.85.2.1 | Head decoration protein D (gpD, major capsid protein D) | 0.485 | BV | *b* |
| b.121.5.1 | Microviridae-like VP | 0.476 | EV | *be* |
| b.121.4.8 | Tetraviridae-like VP | 0.524 | BEV | *be* |
| b.121.2.3 | Major capsid protein vp54 | 0.468 | V | *e* |
| b.121.5.3 | Densovirinae-like VP | 0.468 | V | *e* |
| b.121.4.9 | Birnaviridae-like VP | 0.468 | V | *e* |
| a.62.1.1 | Hepatitis B viral capsid (hbcag) | 0.468 | V | *e* |
| b.121.4.3 | Caliciviridae-like VP | 0.468 | V | e |
| b.121.4.6 | Tymoviridae-like VP | 0.468 | V | e |
| b.121.4.5 | Bromoviridae-like VP | 0.468 | V | e |
| d.254.1.1 | Arterivirus nucleocapsid protein | 0.468 | V | e |
| e.42.1.1 | L-A virus major coat protein | 0.472 | EV | e |
| b.121.5.2 | Parvoviridae-like VP | 0.472 | V | e |
| b.121.7.1 | Satellite viruses | 0.472 | V | e |
| b.121.4.4 | Nodaviridae-like VP | 0.472 | V | e |
| a.190.1.1 | Flavivirus capsid protein C | 0.476 | V | e |
| b.121.4.2 | Comoviridae-like VP | 0.476 | EV | e |
| a.115.1.2 | vp6, the major capsid protein of group A rotavirus | 0.476 | V | e |
| e.28.1.2 | Phytoreovirus core | 0.476 | BV | e |
| a.115.1.3 | Phytoreovirus capsid | 0.476 | V | e |
| b.121.4.1 | Picornaviridae-like VP (VP1, VP2, VP3 and VP4) | 0.481 | BEV | e |
| d.196.1.1 | Outer capsid protein sigma 3 | 0.481 | V | e |
| b.121.6.1 | Papovaviridae-like VP | 0.481 | V | e |
| e.48.1.1 | Major capsid protein VP5 | 0.485 | V | e |
| b.121.4.7 | Tombusviridae-like VP | 0.485 | BEV | e |
| a.115.1.1 | Orbivirus capsid | 0.485 | V | e |
| e.28.1.1 | Orbivirus core | 0.485 | BV | e |
| a.24.5.1 | TMV-like viral coat proteins | 0.489 | EV | e |
| b.121.2.2 | Adenovirus hexon | 0.494 | V | e |
| d.254.1.2 | Coronavirus nucleocapsid protein | 0.494 | V | e |
| a.73.1.1 | Retrovirus capsid protein, N-terminal core domain | 0.506 | EV | e |
| a.28.3.1 | Retrovirus capsid protein C-terminal domain | 0.506 | EV | e |

## Discussion

### Viral and cellular proteomes increase disorder to achieve different goals

A comparative genomic analysis of intrinsic disorder in the proteomes of cellular organisms and viruses revealed a ‘spectrum’ or ‘continuum’ in proteomic sizes, matching results from a previous analysis of nearly 3,500 proteomes with a sampling similar to that of our study [8]. We evaluated this continuum using mean protein length, total number of proteins, and domain use and reuse patterns and their relationship with mean disorder levels of the respective proteomes. This continuum had the smallest and largest proteomes with high intrinsic disorder occupying each end of the spectrum, bracketed by viral and eukaryotic proteomes. Proteomes of archaeal microbes and extremophilic bacteria with highly ordered protein structures occupied the middle of the spectrum, representing the ‘extremes’ of protein interactions [47].

Viruses showed the largest variation of disorder scores in the proteomes we sampled. A tendency of viruses with small single stranded genomes to have proteomes with the highest disorder levels was evident when compared with the bulkier proteomes of dsDNA viruses (Fig. 6c), supporting previous observations [8]. Unlike archaea and bacteria, viruses are enriched in polar amino acid residues with lower hydrophobic amino acid content [47], making their proteins more rigid and prone to low disorder levels. Intrinsic disorder in viruses is mainly employed for RNA binding and interactions with molecules of other organisms [48], as observed by the domain structures listed in Table 3. For example, the nucleocapsid protein of SARS-CoV-2 is the most disordered protein of the coronavirus proteome [49]. The protein holds two nucleotide binding domains, the highly disordered virus-specific N-terminal ‘coronavirus RNA-binding domain’ (b.148.1.1) (Table 3) and the C-terminal ‘coronavirus nucleocapsid protein’ (d.254.1.2), both of which are virus-specific (present in Venn group V) and appeared in the timeline at *nd* =0.494 together with all known virus-specific capsid proteins. The domains are separated by a disordered linker, which together with the two domains hold crucial epitopes for vaccine development that have been actively mutated during the COVID-19 pandemic to change the structure of the molecule and tailor the seasonal behavior of the virus [50]. Disorder increases in viruses are vital for their ability to function and multitask while maintaining compact genomes through reductive evolution [8].

Archaeal proteomes mostly fell in the realm of ‘ordered’ structures, with the exception of Halobacteria. Archaea leveraged disorder in biological functions related to ‘Information’ functions (Fig. S1), mainly for translation [10, 48]. The high disorder content of halophilic archaea is likely due to limitations in prediction methods as disorder estimation methods are developed using non-halophilic proteins under normal saline conditions [8]. The structure and function of proteins harbored by halophilic archaea rely on the presence of salt. Halophilic archaea employ a “salt-in” method to adapt to the presence of salts, thereby maintaining ordered structural conformations at higher salt concentrations [51]. Moreover, the proteomes of halophiles tend to be highly acidic and their proteins are found to be unstable at low salt concentrations.

Most bacterial organisms we analyzed had ordered proteomes, with a fraction of them being moderately disordered. Like archaeal microbes, bacteria utilize disorder in ‘Information’ functions, such as transcription and translation (Fig. S2). Other functions where bacteria leverage disorder include metabolic and catabolic functions, RNA splicing, and pathogenesis (Table 3). Disordered proteins in bacteria are localized in specific cellular components, including nucleosomes, ribosomes, transcription factor complexes, cell walls and flagella [48].

In eukaryotes, most proteomes had moderate disorder, except for parasitic protozoa that possessed more disordered regions than their non-parasitic counterparts [8]. Parasitic protozoa and fungi are thought to have high disorder (Fig. 5, Table 2) as an adaptation to the parasitic lifestyle [52]. Disordered regions in eukaryotic proteins possess significantly high disorder that is highly enriched in linker regions, than their prokaryotic counterparts [53]. The eukaryotic linker regions have higher frequencies of serine and proline with lower frequencies of isoleucine, than those of prokaryotes. Moreover, eukaryotes have amino acid compositions similar to viruses, which points to parallel innovation strategies of increased disorder in both groups. Eukaryotic proteomes may opt for high disorder for their defense mechanisms. Conversely, viral proteomes evolve high disorder (Figs. 2A and 12C) to interact with the immune system of the host [54]. This contrasts with bacterioviruses and bacteria, in which the extent of disorder is not the same. Bacterioviruses have higher disorder levels than bacteria (compare Fig. 4a and Fig. 5a), which is in agreement with a previous study [47]. In addition to ‘Information’ related processes, eukaryotes and viruses utilize disorder in ‘Intracellular processes’ (Fig. S3) and are therefore enriched in posttranslational modification sites [48].

### Intrinsic disorder of structural domains is a late evolutionary development

To study the evolution of intrinsic disorder, we traced disorder onto an evolutionary chronology of domains [21]. We found that the most ancient domains were ordered and that disorder in proteins appeared relatively late in our phylogenomic timeline (Figs. 7 and 8). Disorder was therefore a benefit that was acquired later in evolution. This important conclusion is in agreement with a previous phylogenomic tracing of CATH domains that found flexible structures developed late in evolution [55]. In addition, evidence from a comparative study between ancient and young eukaryotic proteins revealed that ancient eukaryotic proteins have less disorder than younger proteins [56, 57]. Furthermore, disorder in younger eukaryotic proteins were found strongly dependent on GC composition and weakly correlated in ancient proteins, suggesting that the relationship between GC composition and disorder weakens with time. In contrast with our results and this evidence, disordered regions in eukaryotes harbored high GC content and it has been argued that early proteins were disordered and became structured gradually with time [11]. While these results appear contradictory, an evolutionary analysis of intrinsic disorder of loop structures, prior molecular states used to build domain structure [58], revealed that loop prototypes (defined by loop geometry and makeup) that were ancient were more disordered than those appearing later in a chronology of loop structures [46, 59]. Thus, disorder appeared earlier in the more granular loops structures than in the emerging domain structures, showing evolutionary constraints are percolating into different levels of structural organization.

In the context of the shared origin of viruses and cells [20, 21, 60], the most ancient domains that were universally shared by viruses and cellular life (the ABEV Venn group) and appeared at the start of the chronology of domains were ordered and were soon followed by domains with moderate disorder (Fig. 8). This same pattern of an early start of ordered domains was observed with ancient domains shared by cellular life (ABE group) and domains belonging to the Venn groups that followed. Remarkably, the time of origin of the first domain family with high disorder (b.4.1.1) occurred well after the appearance of the BEV group at *nd* = 0.214 and during Phase II (birth of archaeal ancestors)(Table 3), once the stem lines leading to viruses and cells were well established [46]. These early origins support the abundance of ordered proteins in ancestors of both viruses and cells and the rare appearance of domain structures with significant but low levels of disorder (mean disorder of 0.317-0.441) in the emerging stem lines of that time (*nd* bin 0.3 with *nd* values of 0.2-0.3)(Table 3). The strength of disorder levels of the high disorder domains increased in evolution, showing the evolutionary benefits of intrinsic disorder being increasingly adopted by domain structure (Fig. 7e).

When we focused on domains unique or shared between archaeoviruses, bacterioviruses, and eukaryoviruses, i.e. viral domains grouped by the superkingdoms of their hosts, we found genomic distribution, disorder type and history (Fig. 12) largely matched that of our global analysis (Fig. 8). The most ancient domains of the core *abe* Venn group, which is comprised of ordered and moderate disorder structures, were likely acquired by ancient viruses while interacting (or being part of) ancient cells. During that time, the range of existing hosts was limited to common ancestors of life arising during the first two evolutionary phases of the timeline. The higher mean disorder levels and wide distribution of Venn groups *be, e, b* and *ab* suggest viruses may have diversified their disorder repertoire by recruitment of domains with higher disorder levels later in evolution in order to infect a wide range of hosts.

### Intrinsic disorder tailors proteome makeup

The spectrum of disorder versus proteome makeup (proxy for organismal complexity) illustrated in the ‘butterfly-like’ plots of proteomic use and reuse of domains (Fig. 2b) captures a dichotomy in the evolutionary trajectories opted by the ancient ancestors of viruses (proto-virocells) and the ancestors of cells. The dichotomy was also captured by plots of mean disorder against protein number and mean protein length (Fig. 1). While proto-virocells evolved their proteomes via genomic reduction into modern viruses [20, 21], our analysis reveals that viral proteomes gained disorder at the cost of reduced genomic makeup, while at the same time perfecting the ability to integrate with host systems for joint evolutionary success [61]. This trajectory was particularly exploited by RNA and single stranded DNA viruses (Fig. 6c). Most likely, disorder conferred mutational robustness to the otherwise small and vulnerable proteomes of viruses [62]. Protein disorder allowed both expanding interactions with a range of cell targets [63] and functional promiscuity [64]. This helped undermine host immune systems [65, 66] for virus-host integration, thereby aiding in economy of their genomes. The second evolutionary trajectory captured by the ‘butterfly-like’ plots embodies both genomic expansion in cells [67] and cellular diversification [68], which was evident in the proteomes of eukaryotes and particularly in the growing lineages of Metazoa (Fig. 5b). This likely unfolded due to massive co-option of disordered regions that were being developed in protein loops [59] and combined in domains [58], which afforded eukaryotes the ability to evolve advanced functionality by virtue of their large genomic sizes. This cooption is indirectly linked to the remarkable finding of our study that there is a clear increase of mean disorder with decreases in the average length of domains (Fig. 7f), a tendency that is also followed by viral proteins in proteomes (Fig.1b). This relationship is particularly significant because the organization of domains in proteins obeys Menzerath-Altmann’s law of language [69]. Domain length consistently decreases with increasing numbers of domains in proteins showing a tendency of parts to decrease their size when systems enlarge. This tendency towards economy for example manifested in Metazoa, which achieves maximum flexibility and robustness by harboring compact molecules and complex domain organization [69], elements that push in our study towards increasing disorder.

To reconcile the widespread distribution of ordered domains in cells and viruses with our findings of high disorder being mostly exploited by viral proteomes, we tested if viruses recruited highly disordered domains from cells, or vice-versa, they acted as ‘melting pots’ of cellular innovation to aid in host adaptations [61]. Reconciliation involves two alternative explanations, cell-to-virus transfer of disorder domains or virus-to-cell transfer of those innovations to expand host range and adapt to the growing flexibility being developed by their hosts. Comparative proteomic analyses revealed that domains shared with viruses spread more widely (have higher *f*-values) in the proteomes of organisms belonging to Archaea, Bacteria and Eukarya [20, 21, 61]. These results suggest viruses foster innovation and help spread molecular wealth in the proteomes of organisms in the three superkingdoms. Figs. 10 and 11 confirm these results. The significant difference in means of distribution between ordered domains shared in cells and those shared with cells and viruses shows that viruses did not necessarily acquire ordered domains from cells (Fig. 10a). Instead, it is more likely that viruses generated novelties that were then spread to their cellular hosts (see Table 5 for an illustration with capsid domains). It also points to the possibility that ancient viruses may have introduced some of the domains while interacting with ancient cells. Remarkably, no significant differences were detected between the means of distribution of domains with moderate disorder and high disorder shared with cells and those shared between viruses and cells (Fig. 10b and c). Consequently, inferences about the direction of gene-transfer of moderate disorder and high disorder domains, which are evolutionarily more derived, cannot be drawn at this time. We speculate that high levels of disorder evolved in parallel and late in both viruses and cells, employing a similar strategy to maximize disorder while catering to different goals (e.g. ‘confuse-hide-mimic-hijack’ hosts or ‘find-recognize-respond-kill’ the pathogens [70]).

We end by noting that the methodology we present in this paper provides a unique perspective capable of dissecting both evolution of disorder at a scale of billions of years of evolution and an exploration of the relationship between layers of organization in biological systems, i.e. the building blocks versus the wholes (protein loops and domains, domains and proteomes). Disordered regions tend to have high mutation rates and therefore, protein structure provides a far more reliable means of tracing the history of intrinsic disorder.

## Data availability

The data that supports the findings of this study are publicly available in SCOP (https://scop.mrc-lmb.cam.ac.uk) and SCOPe (https://scop.berkeley.edu) repositories. Other data and information supporting the findings of this study are available within the article and its supplementary information files.

## Author contributions

Conceptualization, F.M. and G.C.-A.; methodology, F.M. and G.C.-A.; validation, F.M.; formal analysis, F.M.; investigation, F.M. and G.C.-A.; data curation, F.M.; writing—original draft preparation, F.M.; writing—review and editing, F.M. and G.C.-A.; visualization, F.M. and G.C.-A.; supervision, G.C.-A.; project administration, G.C.-A.; funding acquisition, G.C.-A. All authors have read and agreed to the published version of the manuscript.

