## Supplementary information for "Evolution of intrinsic disorder in the structural domains of viral and cellular proteomes"

SCIENTIFIC REPORTS

The PDF file includes:

Fig. S1. Spread of domains categorized by molecular function and degree of disorder in archaeal proteomes.
Fig. S2. Spread of domains categorized by molecular function and degree of disorder in bacterial proteomes.
Fig. S3. Spread of domains categorized by molecular function and degree of disorder in eukaryotic proteomes.

### Supplementary Figures

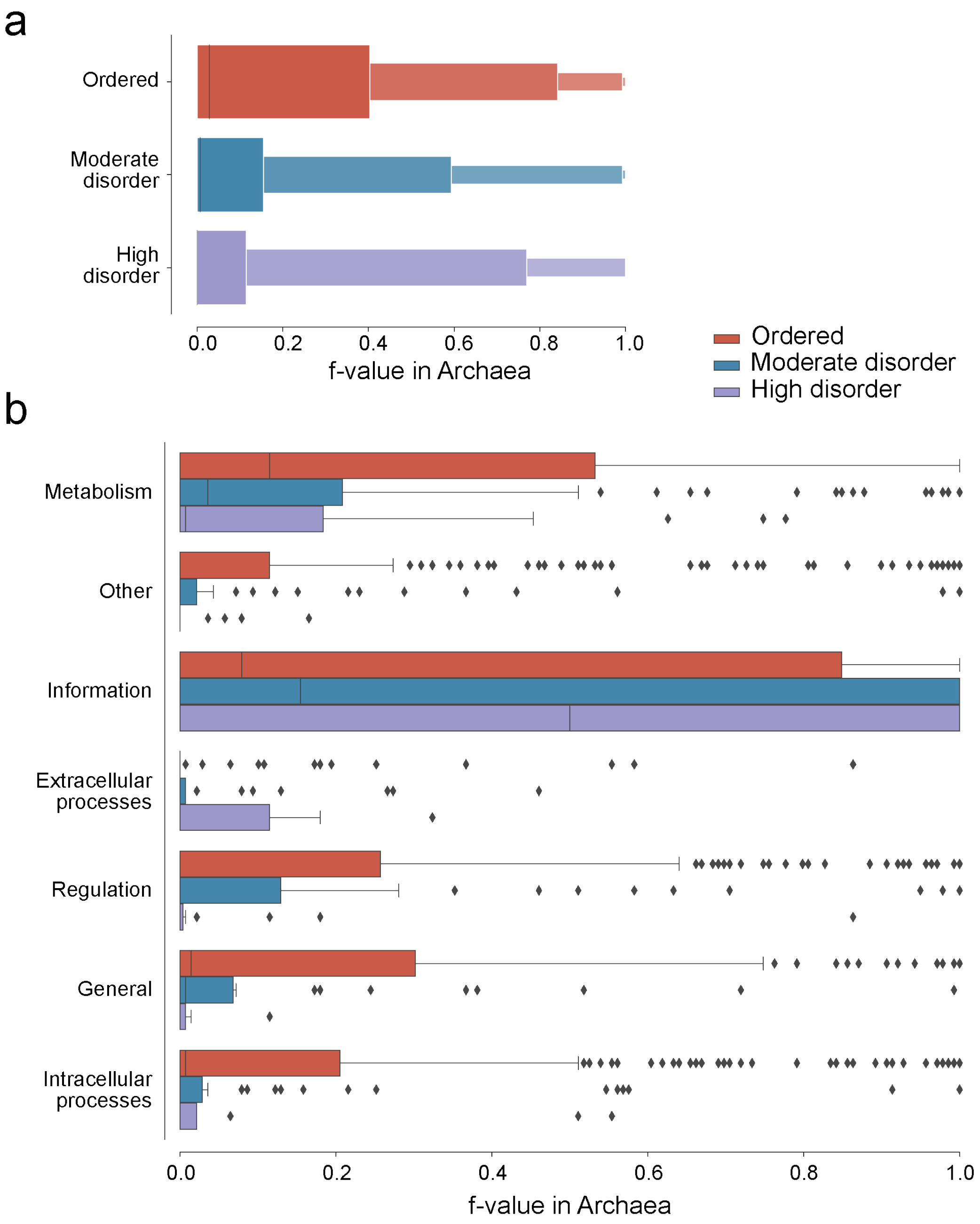


**Figure S1.** Spread of domains categorized by molecular function and degree of disorder in archaeal proteomes.

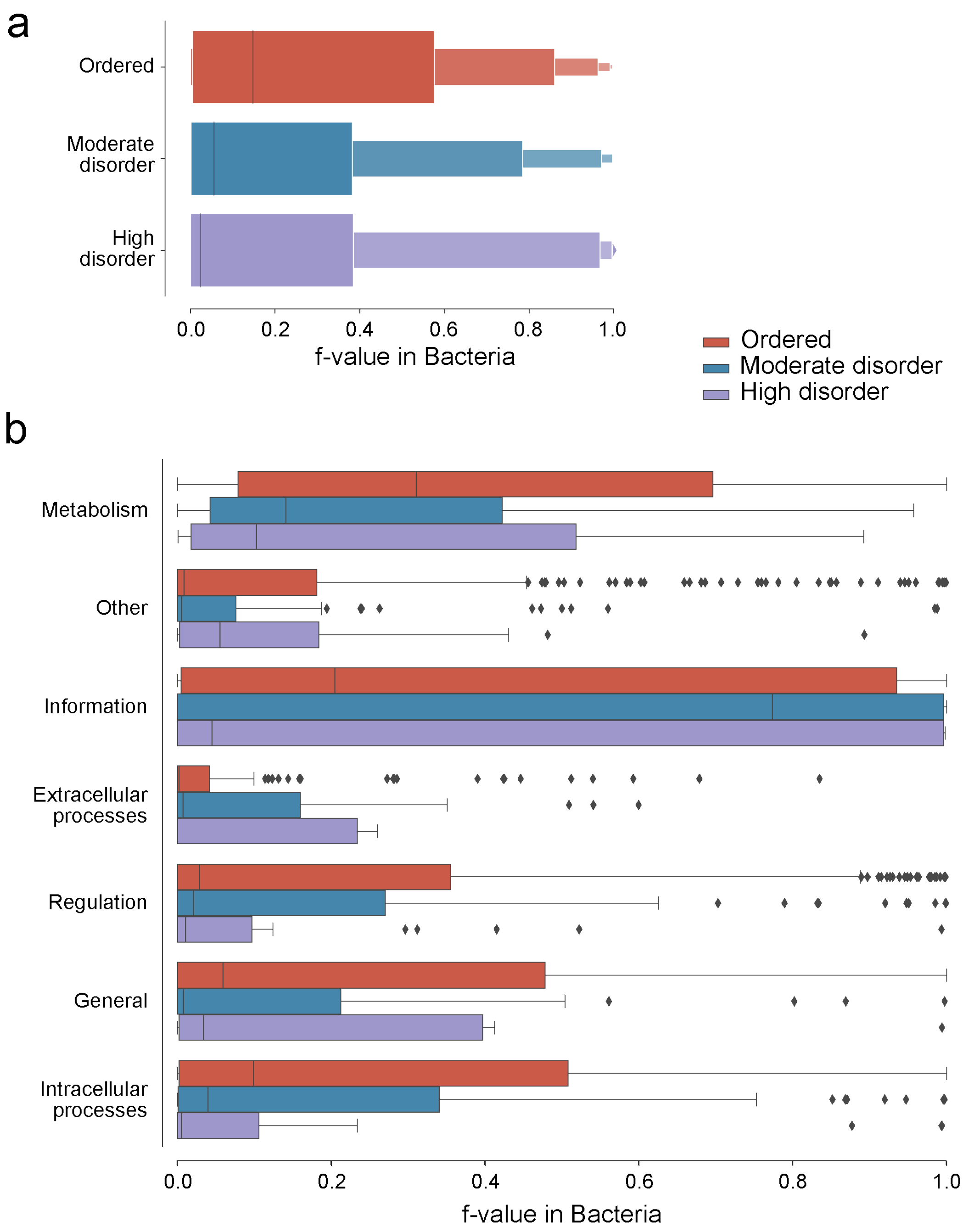


**Figure S2.** Spread of domains categorized by molecular function and degree of disorder in bacterial proteomes.

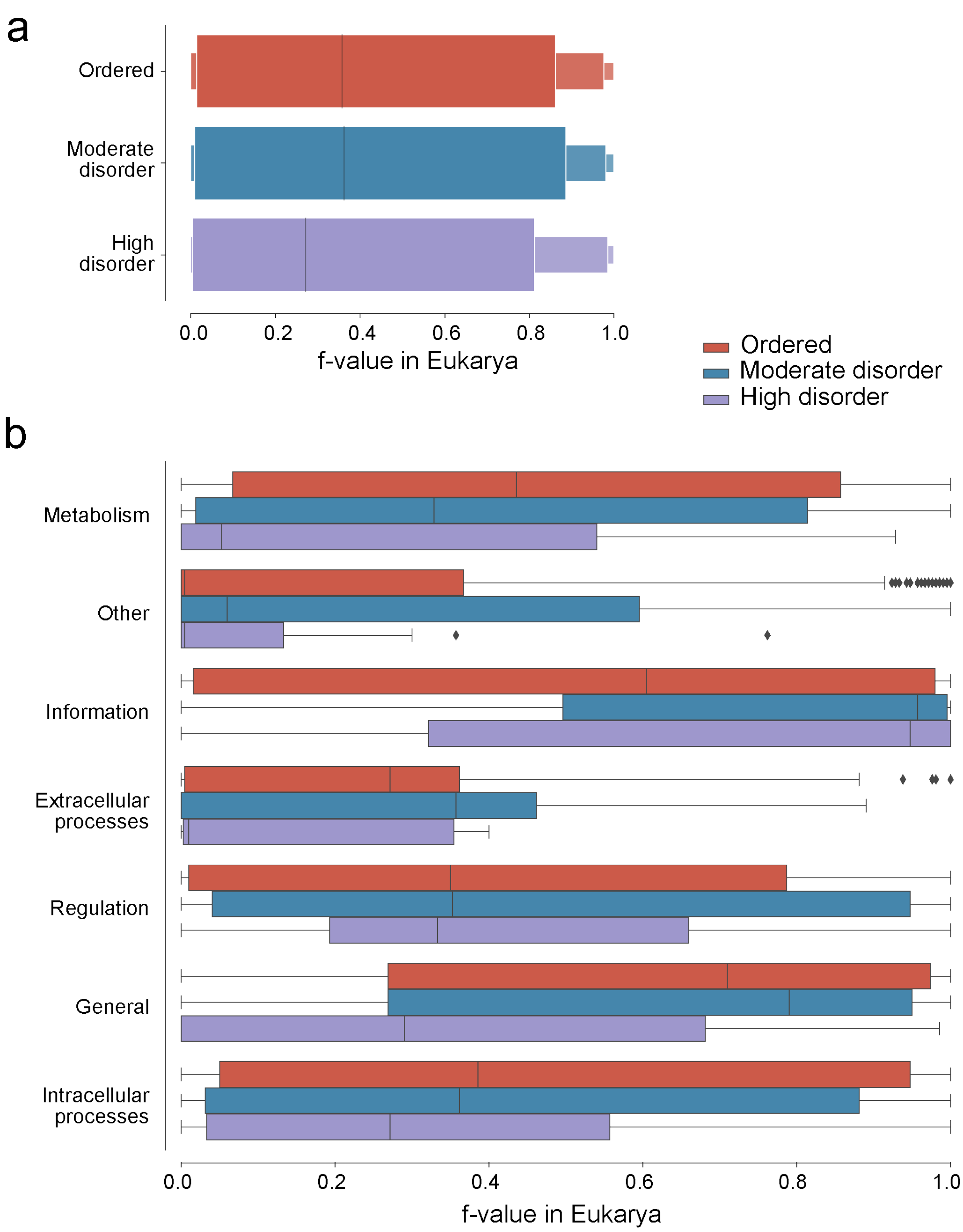


**Figure S3.** Spread of domains categorized by molecular function and degree of disorder in eukaryotic proteomes.
